# Cortical microinfarcts caused by single penetrating vessel occlusion lead to widespread reorganization across the entire brain in a CX3CR1 dependent manner

**DOI:** 10.1101/437509

**Authors:** Alisa Lubart, Amit Benbenishty, Hagai Har-Gil, Hadas Laufer, Amos Gedalyahu, Yaniv Assaf, Pablo Blinder

## 1. Introduction

Dementia, and the associated cognitive impairment that goes along with it, have been attributed to a plethora of different mechanisms. As proper brain function is tightly coupled with the on-demand supply of oxygen and nutrients to active areas, perturbation to blood flow and the resulting neurovascular uncoupling play a crucial role in the ethology of these pathologies [1, 2, 3], even before the onset of the clinical symptoms [4]. Although research has traditionally focused on systemic and large-scale perturbations such as hypertension and stroke, cortical microinfarcts emerge nowadays as a novel and important risk-factor in humans [5, 6, 7], almost doubling the odds for developing dementia as reported in a recent meta-study [8]. Microinfarcts result from the occlusion of relatively small vessels and leave a well defined and restricted histological fingerprint with necrotic tissue extending into less than a cubic millimeter. Despite novel knowledge pointing to a larger proximal area beyond the ishcemic core and penumbra [9], it is still largely unclear how such highly-localized damage can contribute to cognitive decline where affected areas and functions are distributed across different brain areas.

Experimentally-generated single penetrating vessel occlusions can be regarded as the minimally rele-15 vant vascular insult as those lead to the formation of cortical microinfarcts resembling in size an appearance the clinical findings [10, 9]. In turn, occlusions of either pial [11, 12] or capillary vessels [13] do not lead to an ischemic event given the redundancy present on these portions of the cortical vascular network [14, 12]. To asses the potential long-range impact of cortical microinfarcts, we combined here targeted photothrombotic occlusion of single penetrating cortical vessels with diffusion tensor imaging (DTI), tractography and histology. Cortical microinfarcts led to remodeling across large brain regions and to the formation of a microglia/macrophage migratory trail along the subcortical white matter tracts; processes that where dependent on the activity of the microglia/macrophage fractalkine receptor CX3CR1. Combined, we provide here evidence to link local microinfarcts with brain-wide remodeling pointing at microglia/macrophage motility and reactivity as mediators of this phenomena.

## 2. Results

### 2.1. Microinfarcts results in brain-wide structural reorganization

We first set to test whether a single cortical penetrating vessel occlusion could have an effect beyond the expected limits [9]. Since we wanted to capture morphological changes through time and over the entire brain, we turned to Diffusion-Tensor Imaging (DTI) which has been used to observe structural cerebral changes in human and small animals studies following stroke [15, 16]. Here, DTI allowed us 31 to longitudinally track remodeling over time and compare it to a baseline conditions of each animal prior to the occlusion. Anesthetized adult mice underwent targeted photothrombotic occlusion of single penetrating-vessel (Figure 1A) which resulted in the formation of a cortical microinfarct (Figure 1B). We chose to perform the occlusion in animals implanted with a polished and reinforced window (PoRTs) as this particular preparation enabled imaging and induction of the photothrombotic occlusion while minimizing or completely avoiding inflammatory response [17, 18] which can bear potential confounding effects.

**Figure 1:**
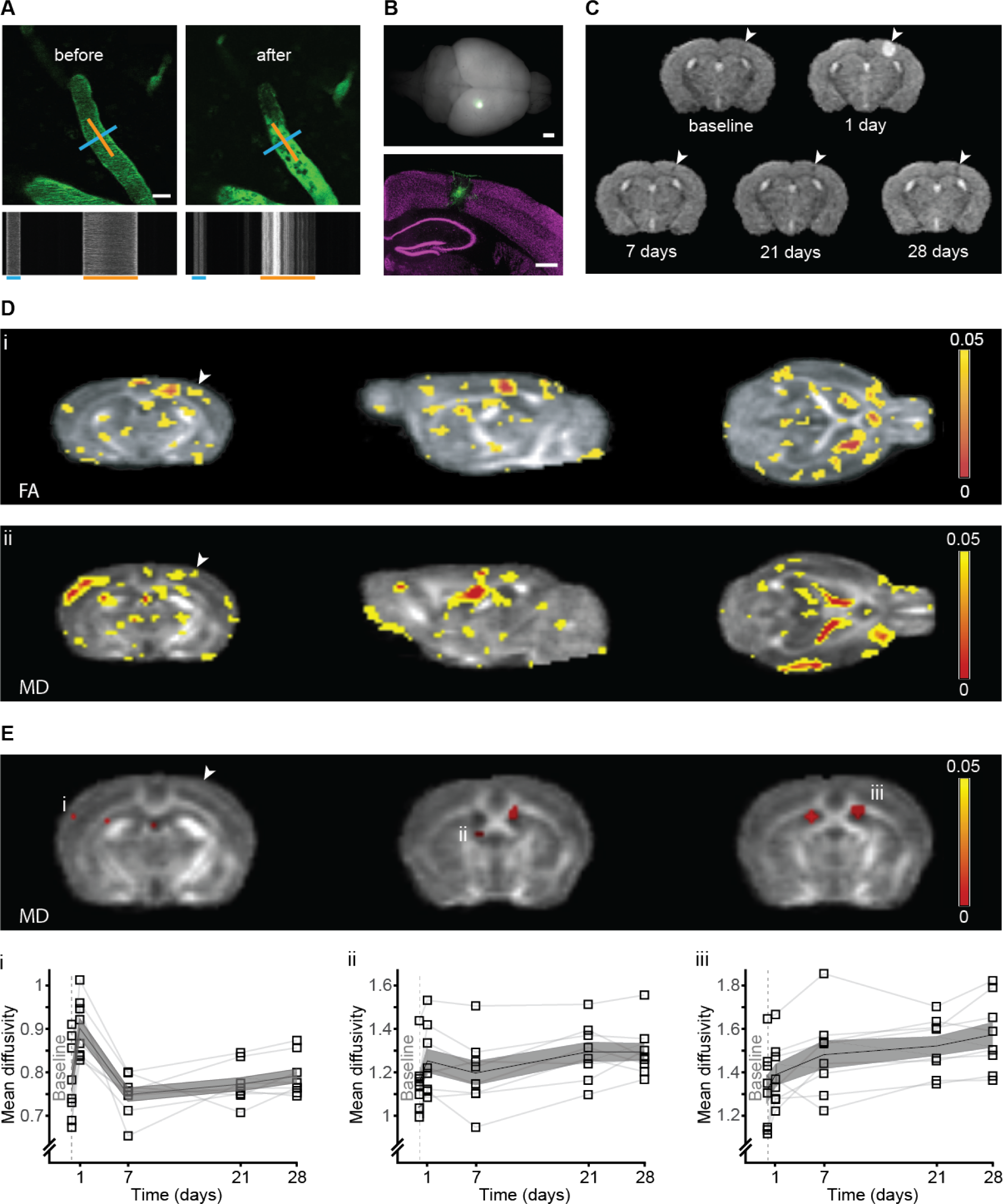
(Previous page) Targeted photothrombotic occlusion of single penetrating vessel leads to widespread structural remodeling. (A) Intravital imaging of a C57BL/6J WT mouse pial vessels before (left) and 5 min after (right) occlusion of a single penetrating artery. Blood plasma was labeled with FITC-dextran (2000kDa). Line-scan of the blood flow pattern in the targeted blood vessels allowed extraction of the blood vessel diameter and direction of the blood flow; blue and orange lines respectively and corresponding time-space linescans (bottom). (B) A whole mouse brain 14 days after targeted photothrombotic occlusion, note the FITC-dextran retention surrounding the occluded area (green, bottom panel). NeuN staining (magenta) indicated the occlusion size in the cortex. (C) Temporal evolution of T2 in a single WT mouse: hyperintense signal at the infarct area 24h post occlusion disappears at 7 days post occlusion until only a small hypointense signal at the same area was left at 28 days post occlusion. (D) Statistical FA (i) and MD (ii) maps of the main effect of time post micro-occlusion superimposed on an averaged FA map of all mice. Voxels that exceeded a statistical threshold of p<0.05 (Repeated measure ANOVA, n=8) are colored according to the threshold they exceeded. (E) Clusters that exceeded a statistical threshold of p<0.05 corrected for multiple comparisons (FDR) of the MD maps. Examples of clusters showing significant change in MD through time, in the contralateral somatosensory cortex (i), contralateral fimbria of the hippocampus (ii) and and in the ipsi and contralateral ventricles (iii). (D-E) Arrowhead indicates the occlusion area. Scale bar: A:20m, B: 1mm, 500m.

Nine wild type C57BL/J mice were scanned before micro-occlusion induction (baseline), and 1, 7, 21 and 28 days following micro-occlusion (Figure 1C). T2-weighted imaging 1 day post micro-occlusion revealed a hyper-intense signal at the lesion site, indicating tissue damage and edema formation. This hyper-intense signal extended from the pial surface into the cortical gray matter, and in some mice reached the underlying white matter of the corpus callosum. The hyperintense signal was lower in the 1-week time point, indicating reduction in edema through time, while at 4 weeks a small hypointense signal was visible at the lesion site (Figure 1C, arrow), indicating gliosis formation ([19]).

Tissue remodeling was probed by measuring and analyzing two parameters obtained from DTI: mean diffusivity (MD) and fractional anisotropy (FA). MD measures the average magnitude of the molecular motion while FA measures its directionality [20]. Initially, we analyzed MD and FA changes over time using a region based analysis of the infarct area of each mouse and the corresponding area in the contralateral hemisphere. As expected ([21, 22]) regional MD and FA values in the infarct area decreased one day following occlusion and gradually increased thereafter (Supplementary Figure S1A i+ii) while in the exact same region on the contralateral hemisphere there were almost no changes following occlusion throughout time in both parameters (Supplementary Figure S1B i+ii). Looking beyond these area revealed extensive remodeling. A voxel-based repeated measure ANOVA (five time points), highlighted several clusters where a significant (p<0.05, ANOVA) change in FA (Figure 1D, i) and MD (Figure 1D ii,) values occurred through time, before correction for multiple comparisons ([23]). Significant changes after correction for multiple comparisons (p<0.05) in regional mean diffusivity through time appeared 57 in brain regions located several millimeters away from the occlusion site, including -among others– regions such as the contralateral somatosensory cortex (Figure 1Ei) and the contralateral fimbria of the hippocampus (Figure 1Eii). Similarly, both ipsi- and contralateral ventricles showed remodeling (Figure 1Eiii). These results indicate that a microinfarcts lead to changes in tissue structure far from the local lesion site affecting areas across the entire brain.

### 2.2. Occlusion of a single penetrating-vessel in the somatosensory cortex results in distant accumulation of FITC-dextran along subcortical white matter

As part of the targeted-phototrombotic occlusion protocol, a 2000kDa dextran conjugated to fluorescein was injected into the blood stream to visualize blood vessels, and measure flow and vessel diameter [11]. Accumulation of the dextran around the ischemic area is commonly observed [11, 24, 25, 26]. To 67 our surprise, histological examination of brains of mice after their last DTI scan, 28 days following micro occlusion induction, revealed accumulation of FITC-dextran well beyond the occlusion site, spreading along the sublesional white matter (Figure 2A) and to a lesser extent, over the pial meninges (Supplementary Figure S3). In what follows, we focused on the white matter phenomena which termed as trail. To this end, separate animals were induced with a single micro-occlusion and sacrificed at different time points. The trail spread over the subcortical axonal tracts, and could be clearly observed in sections anterior and posterior to the area of infarct. In fact, in about 15% of the cases it crossed to the contralateral hemisphere (Figure 2B). If the trail reflected some pathological process or remodeling of the white matter, this could turn to be the missing link between local microinfarcts and their long range effects identified by DTI (Figure 1), prompting us to further explore this phenomena.

**Figure 2:**
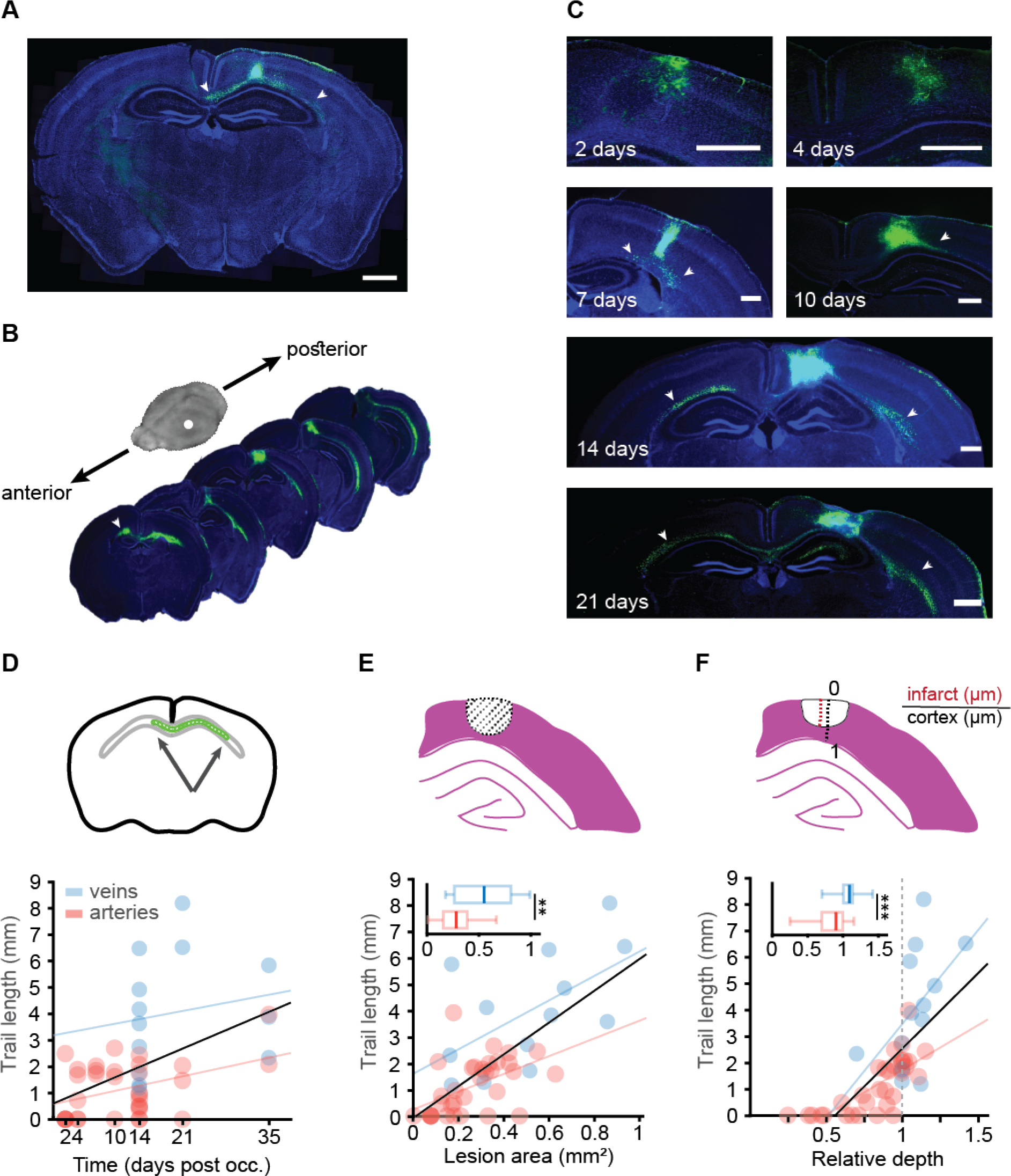
(Previous page) Occlusion of a single penetrating blood vessel leads to distal FITC-dextran accumulation. (A) Coronal slice of DTI-scanned mouse (Fig 1C) shows FITC-dextran retention in the occlusion site and the underlying white matter (green). (B) Representative brain slices of a mouse sacrificed 14 days after the occlusion, demonstrates FITC presence at more anterior and posterior areas of the white matter from the occlusion site. (C) FITC-dextran presence at different time points after targeted photothrombotic occlusion induction: 2, 4, 7, 10, 14, 21 and 35 days. Cell nuclei marked with DAPI. At early time points the FITC-dextran was at the cyst area but later reached the subcortical white matter, creating a trail of FITC labeled cells that even crossed to the contralateral hemisphere. (D) Trail length was correlated with time post occlusion for all blood vessels (p=0.001, R^2^=0.22, Black), but separately it was significantly correlated with arteries (p=0.0195, R^2^=0.1637, red) but not for veins (p=0.56, R^2^=0.03, blue). (E) Infarct area was correlated with trail length for all blood vessels (p<0.0001, R^2^=0.4814, Black), and separately for arteries (p=0.0038, R^2^=0.24, red, n=33) and veins p=0.037, R^2^=0.3385, blue, n=13). Total lesion area was larger in veins (p=0.0057, Mann-Whitney). (F) Infarct relative depth was also correlated with trail length for all blood vessels (p<0.0001, R^2^=0.488, Black), and separately for arteries (p<0.0001, R^2^=0.5536, red, n=33), but not for veins (p=0.0705, R^2^=0.2671, blue, n=13). Total infarct depth was larger in veins (p=0.0007, student’s t-test). Insets in figures D-F show the metric that was used to measure each variable. Scale bar: A:1mm, C:500m.

As BBB integrity is impaired in the core and penumbra areas ([27]), we hypothesized that the source of the FITC is nearby the occluded vessel as opposed to potentially damaged vessels along the white matter. To distinguish between these two alternatives, we used a red dextran (TexasRed, 70kDa) for visualizing blood vessels when performing the photothrombotic occlusion, and injected the green FITC-dextran either immediately after (i.e. day 0) or in subsequent days post occlusion (days 1,2,7 and 14). All animals were sacrificed 14 days following occlusion. We only observed the formation of a trail for mice injected at day 0. In all other time points, the FITC-dextran accumulated in the infarct site where the BBB was impaired ([26]), but was not detected along the white matter nor pial surface (Supplementary Figure S2), indicating that the FITC did not infiltrate from the periphery several days after the occlusion, but rather from the infarct site, either immediately after the clot formation or in the course of the following 24 hours.

To investigate the temporal dynamics of the trail formation, mice were sacrificed at different time points (namely, 2, 4, 7, 10, 14 21 and 35 days) following targeted photothrombotic occlusion of either a vein or an artery of similar diameter (p=0.5887, Student’s t) (Supplementary Figure S4). In the first week following occlusion, the FITC was visible mainly in the infarct area (days 2 and 4). However, from day 7 to 35 post-occlusion, a trail formed along the corpus callosum and to the internal capsule (Figure 2C). Fourteen days after the occlusion and later, the trail reached the contralateral hemisphere along the corpus callosum (Figure 2B). Trail length was positively correlated with time post-occlusion when considering all occluded vessel types together (r=0.47, p=0.001) or penetrating arteries (r=0.4, p=0.019), but not veins alone (r=0.17, p=0.56) (Figure 2 D). Area and depth were not correlated with time following occlusion in neither vessel types (for area: arteries - r=0.155, p=0.385 and veins - r=0.25, p=0.41; for depth: arteries - r=0.288, p=0.103 and veins - r=0.367, p=0.217). In addition, we detected a strong correlation between trail length and lesioned area cross section (r=0.69, p<0.0001) and depth into the cortex (r=0.7, p<0.0001) (figure 2 E-F). Notably, occlusion of veins resulted in larger (p=0.0057, Mann-Whitney) and deeper (p=0.0007, student’s t-test) lesions (Figure 2E,F). Thirty-two percent of all lesions reached the white matter (15 of 46), while most of these were veins (9 of the 15).

### 2.3. Microglia/macrophages migrate along the corpus callosum following penetrating vessel occlusion

To further understand the nature of the observed trail, we performed a immuno-histological survey to identify cells that uptake the FITC-dextran. We chose the 14 days post-occlusion time point from the previous experiments for the analysis, as at this time a clear trail was observed along the corpus callosum in most animals. Consistent with previous reports ([28, 29, 30]), we observed microglia/macrophages and astrogliosis surrounding the lesion site. While microglia/macrophages occupied the infarct, astrocytes did not infiltrate the lesion core, forming a glial scar, as previously reported ([31]) (Supplementary Figure S5). Notably, microglia/macrophages in the penumbra were also FITC-positive, while astrocytes were not co-localized with FITC (Supplementary Figure S6).

Importantly, the vast majority of the FITC-dextran outside the infarct itself was found to be inside cells, specifically microglia/macrophages (83% positive for both Iba1 and FITC). In line with this finding, 51% of the FITC-positive cells were also positive for CD45 which is preferentially expressed by macrophages and activated microglia (as opposed to Iba1 which is constitutively expressed). In contrast, FITC was rarely found inside astrocytes (2.6% positive for both GFAP and FITC). Doublecortin (DCX), a marker for newly formed neurons, was also not colocalized with the FITC-dextran (¡2%). Eight percent of the FITC-positive cells were oligodendrocytes (colocalized with Olig2). This last observation was replicated using transgenic mice expressing TdTomato under the control of the NG2 promoter, labeling pericytes and oligodendrocytes precursor cells [32] (OPCs; 6% positive for both TdTomato and FITC; Figure 5 A+B).

To verify if the trail is not formed due to passive dextran leakage but as a result of actual microglia/macrophage spatial reorganization, we performed again single artery occlusion albeit in CX3CR1^GFP/+^ mice, without labeling blood plasma with FITC-dextran. In the CX3CR1-GFP mice either one (heterozy-gous) or two (homozygous) copies of the CX3CR1 allele are replaced with GFP in cells under control of the endogenous CX3CR1 locus (microglia, macrophages, dendritic cells and more). Arteries were labeled with Alexa fluor 633 hydrazide, a specific arterial marker [33] with good spectral separation from the GFP expressed under CX3CR1 [34] (Figure 5C left). A photothrombotic occlusion was induced, resulting in a small infarct (Figure 5C right). In line with our previous experiments, fourteen days following occlusion of a single penetrating artery, a trail of CX3CR1+ cells formed along the corpus callosum. In 131 general, expression levels of CX3CR1 are higher in activated microglia, leading to increased GFP expression ([35, 36]). Therefor, to distinguish between migration and local activation, we measured GFP+ cell density in 5 regions of interest (ROIs) along the corpus callosum of CX3CR1^GFP/+^ mice 14 days after photothrombotic occlusion, and compared it to control mice (Figure 5D). For each ROI, we computed the distance along the white matter tracts to the center of the infarct area (or an estimated center for 136 corresponding location in the control group). We found a significant difference in the microglia distribution between groups as a function of distance from the infarct (p<0.01 for the group difference, p<0.001 for the distance effect and p<0.01 for the interaction between group and distance, mixed-effect general linear model). In control mice, resident microglia displayed uniform density throughout the different ROIs along the subcortical white matter. In contrast, following targeted occlusion, GFP+ cell density 141 increased near the infarct area and decreased with distance from the infarct (Figure 5E). This result implies microglia/macrophages increased more pronouncedly their density along the trail rather than their GFP expression level. To verify that the GFP+ cells migrated to the corpus callosum from the lesion core and not from adjacent cortical areas, we measured GFP+ density in the cortex above the corpus callosum in five ROIs selected in similar locations 14 days following occlusion, in CX3CR1^GFP/+^ mice 146 compared to control animals, and found no difference between the groups (p¿0.05) (Supplementary Figure S8). Proliferation of resident microglia has been shown to occur in the surrounding of the ischemic core following photothrombosis [37]. Therefore, to distinguish between migration and proliferation of microglia we tested for the presence of proliferating microglia along the trail. CX3CR1^GFP/+^ mice were injected with BrdU, a nucleotide analog. While BrdU labeling was evident inside and adjacent to the infarct site, there was low double-labeling of GFP+/BrdU+ along the ipsilesional corpus callosum, and no double labeling in the contralesional hemisphere (Supplementary Figure S7). These results indicate that the trail consists of resident albeit migratory microglia/macrophages.

### 2.4. CX3CR1 pathway mediates trail length following arterial occlusion

CX3CR1 signaling pathway plays a key role in microglia/macrophage migration and activation ([38, 39]). To test whether trail formation is mediated by this pathway, we compared the impact of a single penetrating artery occlusion between CX3CR1^GFP/GFP^ (i.e. CX3CR1 knocked-out) and heterozygous (HET) mice, fourteen days following occlusion (Figure 4A). Notably, the average diameter of occluded arteries was similar in both groups (p=0.788, Student’s t-test) (Supplementary Figure S9A-B) and infarct areas did not differ between knock-out (KO) and heterozygous animals (p=0.387, Student’s t-test) (Figure 4B). Although GFP+ cells accumulated in the infarct core of both genotypes, the mean trail length 162 was significantly higher in CX3CR1^GFP/+^ compared to the CX3CR1^GFP/GFP^ mice (p=0.0047, MannWhitney) (Figure 4C). Next, we tested the correlation between lesion area and trail length and infarct depth with trail length for both genotypes. Area (r=0.57,p=0.017) and depth (r=0.63, p=0.007) were positively correlated with trail length in heterozygous mice but not in KOs (r=0.42, p=0.21 and r=0.3, p=0.27, respectively) (Figure 4D-E).

**Figure 3:**
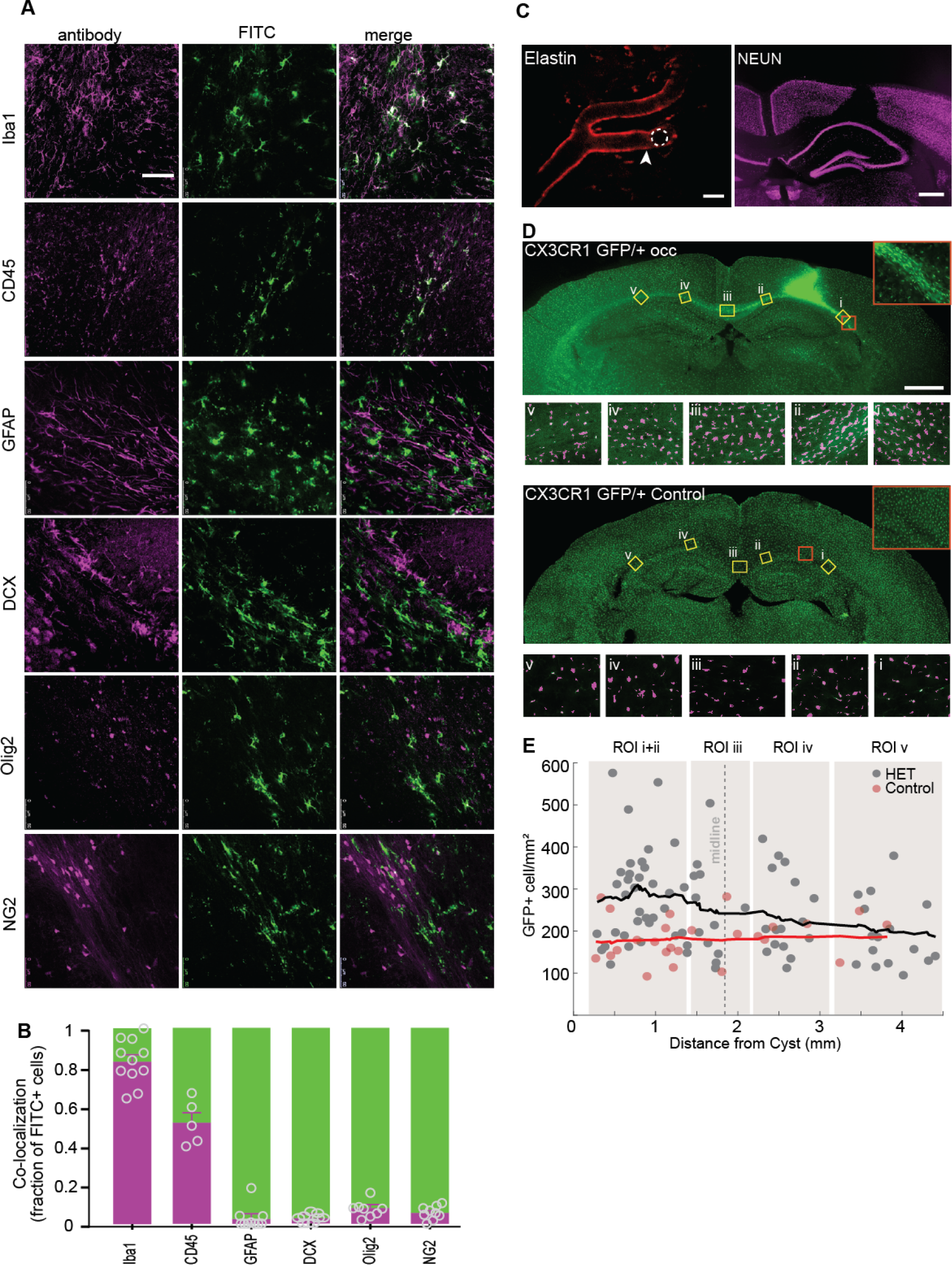
(Previous page) The trail is formed mainly by microglia/macrophage cells. (A) Analysis of brain sections that were obtained from wild type C57BL/6 adult mice, 14 days post targeted photothrombotic occlusion. Sections were stained for different markers in order to characterize the identity of the FITC-dextran binding cells. The majority of FITC-dextran positive cells (green) along the trail that was formed on the corpus callosum were co-labeled (white) with Iba1 (83%) microglia/macrophage marker, and CD45 (51%) leukocyte marker. Astrocytic (GFAP), Oligodendrocytic (Olig2) and neurogenesis (DCX) markers were not colocalized with the FITC-dextran (2% colocalization), whereas NG2 mice revealed 6% co-localization that may be due to their ability to differentiate into microglia cells. (B) Quantification of colocalization of FITC-dextran with the different markers. (C) Two-photon image of a penetrating artery labeled with Alexa fluor 633 hydrazide, arrowhead points to the occluded blood vessel (left). Targeted occlusion in CX3CR1^GFP/+^ mice without labeling the blood plasma led to small infarct in the cortex as visualized by NeuN staining (right). (D) GFP+ density was measured in 5 ROIs along the corpus callosum of CX3CR1^GFP/+^ occluded and control mice, 14 days after micro-occlusion induction. Representative images of automatically identified cells in each ROI are shown below each group (magenta). (E) CX3CR1-GFP cell density plotted as distance from the infarct core, with trend lines (running average, width 25 data points) for distance from infarct core. Mixed-effect general linear model showed that the occlusion led to higher microglia/macrophage density on the white matter tracts near the occlusion site that decreased further away from the occlusion to control levels; all model terms have significant p-values below 0.001, DF = 101. Scale bar: A: 50m, C: 20m, 500m, D: 1mm.

**Figure 4:**
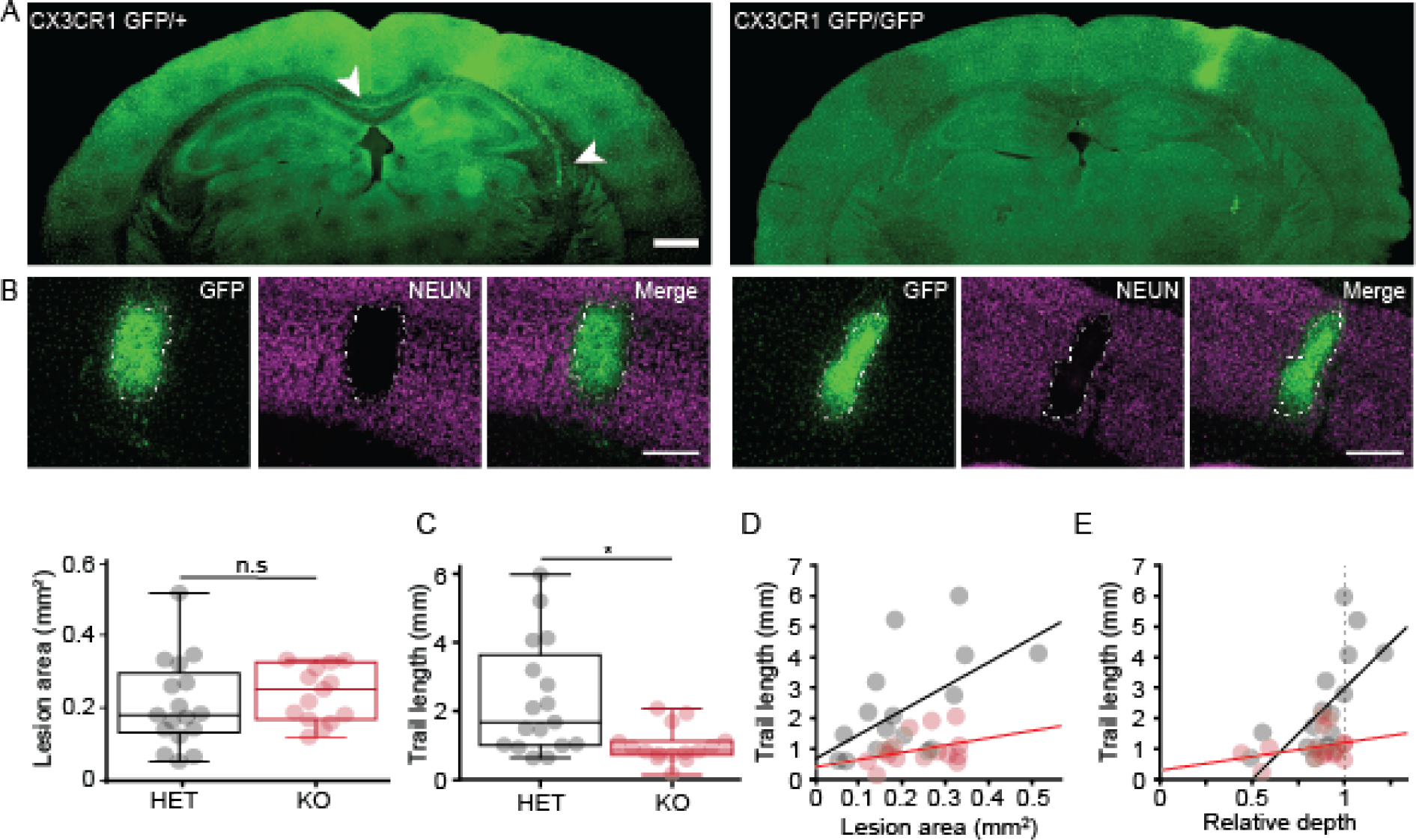
Trail length following targeted photothrombotic occlusion is CX3CR1 dependent. (A) Representative images of CX3CR1 heterozygous and KO mice 14 days after targeted photothrombotic occlusion. (B) The occlusion led to GFP+ cell accumulation in the cyst of both genotypes. The infarct area, delineated in the NeuN staining, was not different between CX3CR1^GFP/+^ (0.210.03mm^2^, n=17) and and CX3CR1^GFP/GFP^ mice (0.240.02 mm^2^, n=15, p=0.387 Students t-test). (C) The mean trail length was significantly longer in CX3CR1^GFP/+^ mice compared to CX3CR1^GFP/GFP^ (p=0.0047, Mann-Whitney), reaching as far as the contralateral hemisphere in 30% of the cases. (D) Correlation between trail length and infarct area for CX3CR1 ^GFP/+^ (R^2^=0.324, p=0.0171, black) and CX3CR1^GFP/GFP^ (R^2^=0.177, p=0.2107, red). (E) Correlation between trail length and relative infarct depth for CX3CR^GFP/+^ (R^2^=0.394, p=0.007, black) and CX3CR1^GFP/GFP^ (R^2^=0.092, p=0.2715, red). Scale bar: A: 1mm, B: 500m.

### 2.5. CX3CR1 pathway mediates structural reorganization following arterial occlusion and determines the impact on white matter tracts

Given the differential response in trail length observed for the different genotypes, we looked for potential remodeling difference across the entire brain using longitudinal DTI imaging in a new set of experiments. A voxel-based mixed-design analysis between group (WT and CX3CR1-KO) and time (5 DTI scans), revealed a significant (p<0.05) main effect for the group factor in several clusters throughout the brain, for MD (Figure 5A) and FA (Figure 5B) maps. The voxel based analysis also revealed signifi-cant multi-regional interaction between group and time in MD values, indicating that in each genotype the occlusion had a different structural impact as a function of the time that has past from the occlusion (Figure 5C). The clusters were located in both the cortex and the white matter, including in the contralateral hemisphere (Figure 5D).

**Figure 5:**
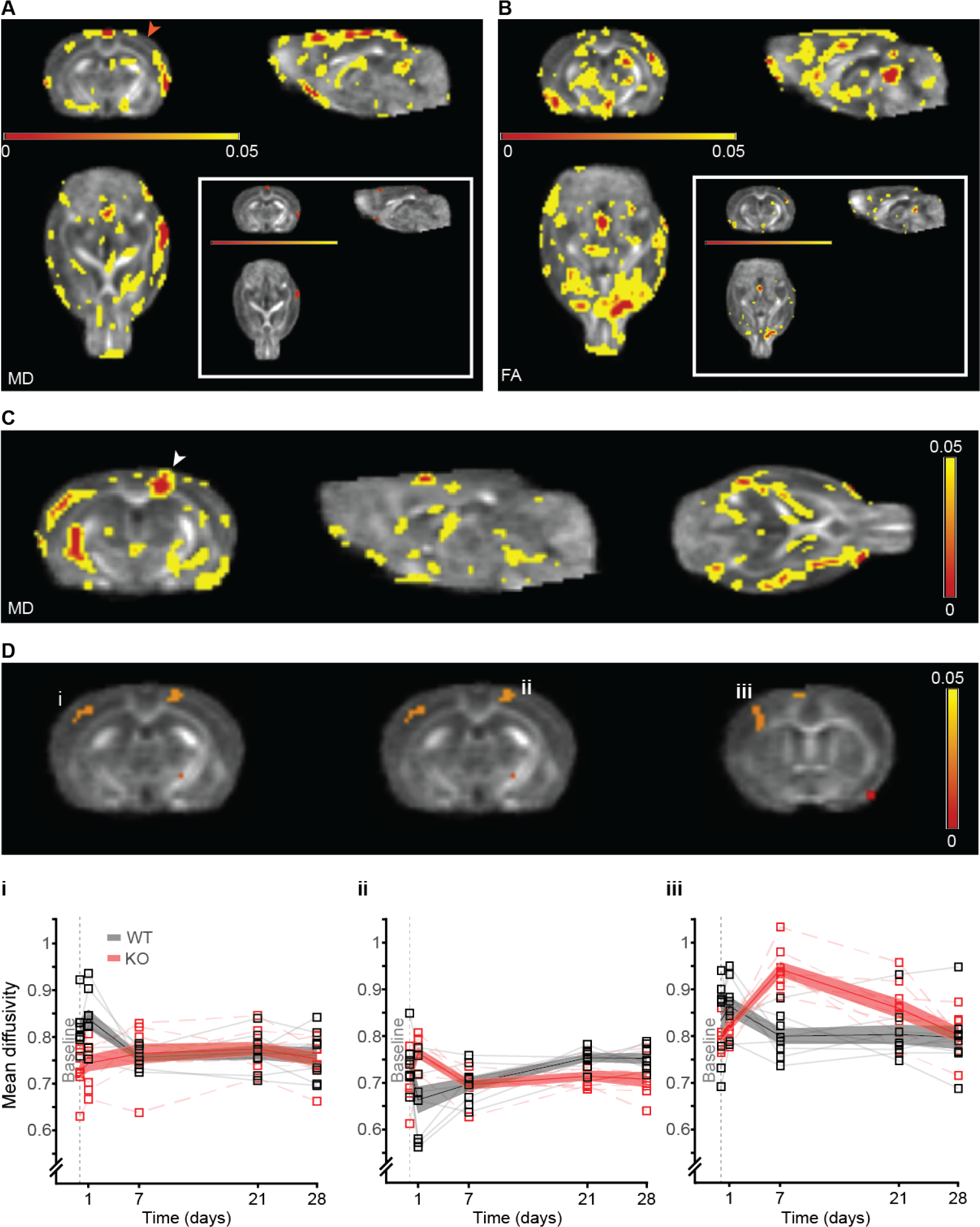
(Previous page) WT and KO respond to the microinfarct in a different manner. (A) Statistical parametric maps of the the main effect of group, regardless of time point, between WT (n=8) and KO (n=8) mice for mean diffusivity (MD) and (B) fractional anisotropy (FA). The statistical maps are superimposed on an averaged FA map of all scanned mice. Voxels that exceeded a statistical threshold of p<0.05 (repeated measure ANOVA) are colored according to the threshold they exceeded. Inset shows regional clusters that exceeded a statistical threshold of p<0.05 corrected for multiple comparisons (FDR). (C) Multiregional interaction between scan time and group (p<0.05, repeated measure ANOVA) (white arrowhead indicates the occlusion area) of the MD. (D) the clusters after FDR correction (p<0.05) for the MD parameter, including contralateral primary somatosensory dorsal area (i), ipsilesional somatosensory cortex (occlusion site) (ii) and contralateral primary somatosensory frontal area (iii), show that the two genotypes respond to the targeted occlusion in a different manner.

To study fiber integrity characteristics following arterial occlusion, we performed post-hoc tractography analysis with corpus callosum as seed region. FA values from the dorsal part of the corpus callosum (splenium) were extracted for each animal (Figure 6A). The mean FA value of each group was plotted throughout the length of the tract. FA values 28 days following occlusion were compared to pre-occlusion baseline. In wild type mice, FA values increased in the contralateral hemisphere, with no change in the ipsilesional hemisphere. In contrast, CX3CR1^GFP/GFP^ mice displayed the opposite behavior, as mean FA values increased in the ipsilesional hemisphere and did not change in the contralateral hemisphere (Figure 6B). This result indicates that CX3CR1 mediates the impact of arterial occlusion on fiber integrity, with distal effects in animals expressing CX3CR1 compared to the local effects in the knock-out mice.

**Figure 6:**
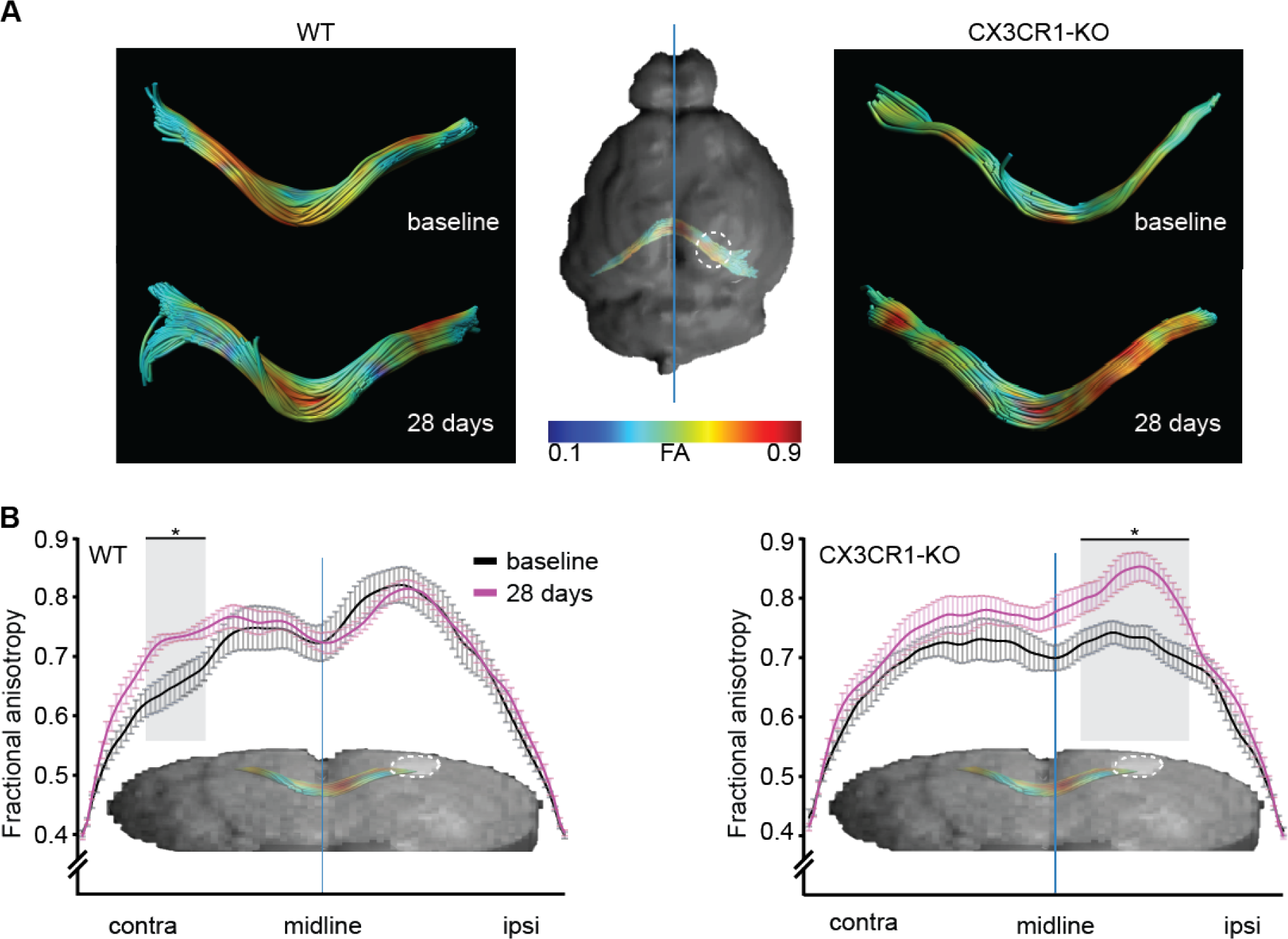
Microinfarct alters fiber-integrity in WT mice in a more distal manner compared to the local effect in CX3CR1^GFP/GFP^ mice. (A) FA values across the corpus callosum (CC) were compared between the last time point (4 weeks post occlusion) to the baseline scan. Representative images of the tracts color coded for FA values, from low (blue) to high (red). Mean FA values were plotted as function of location at baseline scan (black) and 28 days post occlusion (magenta). (B) A significant difference (p<0.005, ANOVA followed by Tukey’s post hoc) in FA values between the two time points was found, along distinct areas of the corpus callosum, such as the ipsilesional CC of CX3CR1^GFP/GFP^ mice and contralesional CC of the WT mice. Asterisk marks the areas where a significant difference between the means of the two time points was found.

To visualize the microglia-myelin interaction we stained sections collected for the histological analyses experiments (14 days following occlusion) for myelin basic protein (MBP), and examined CX3CR1 (green) and myelin (red) co-localization in the corpus callosum. In control mice (i.e. no occlusion) of both heterozygous and KO mice - the myelin staining appeared intact, with myelin sheets across the corpus callosum with only few CX3CR1 cells in the region. However, a single penetrating artery occlusion in CX3CR1 ^GFP/+^ mice resulted in aberrant myelin staining and increased number of GFP+ cells accumulated in demyelinated regions in a chain-like formation. In CX3CR1-KO mice this phenomenon was attenuated - less accumulation of GFP+ cells and the myelin appeared more intact compared to the occluded CX3CR1^GFP/+^ mice (Supplementary Figure S10).

## 3. Discussion

In the current study we report the previously unappreciated long-range structural impact of single cortical microinfarts engendered by occlusion of a single penetrating vessel. Using diffusion tensor imaging (DTI), we show that such restricted occlusions and the resulting microinfarct lead to widespread structural changes in different brain regions, distant from the occluded vessel and the microinfarct, affecting both 201 the ipsilesional and contralesional hemispheres (Figure 1). Moreover, we discovered the formation of a cellular migratory path along the subcortical axonal tracts that forms days to weeks after the vascular insult (Figure 2) and can cross to the contralateral hemisphere through the corpus callosum. The length of this migratory path was found to be dependent on time and size of the infarct (Figure 2). A histological survey allowed us to point at microglia/ macrophages as main components of the migratory path (Figure 5) and provide clear evidence that they migrated along the white matter tracts to form the observed spatial pattern. We further show that this migration was CX3CR1-dependent (Figure 4) and that this molecular pathway also mediates structural reorganization (Figure 5) and modulates axonal fiber integrity following the formation of a cortical microinfarct (Figure 6).

Past work has demonstrated that the histological signature left by cortical microinfarct (generated by the experimental occlusion of single penetrating vessels as done in this work) underestimated the volume of affected tissue. Summers and coworkers [9] elegantly showed that the affected volumes can be up to 12 times larger than the one observed with anti-NeuN staining or with different MRI modalities. The evidence we present here further emphasizes this notion, as we show that while occlusion of a single penetrating vessel results in a small infarct, it leads to structural organization in distant and noncontiguous areas beyond the infarct site and peri-lesion site, affecting also the contralateral hemisphere (Figure 1). Similar extensive remodeling cases have been reported, both in pre-clinical work and in the clinic, for conditions that involved much larger insults such as unilateral strokes [40, 16, 41] and traumatic brain injuries (TBI) [42, 43]. Here, clusters of voxels in different brain areas proximal and distal to the infarct penumbra showed significant changes in MD values as a response to the microinfarct induction. It is well accepted that increased MD values point to reduction in tissue integrity, caused by neuronal death or cyst formation [44]. In turn, decreased MD values might suggest processes such as migration, increase in cell density, and plasticity [45]. We have observed a plethora of MD longitudinal changes associated with different clusters, with both increases and decreases compared to baseline (for e.g. Figure 5Di-iii).

In the current study we observed a dextran-trail that originated from the occlusion core and spread throughout the sublesional white matter. This finding was somewhat surprising, as other studies that used the exact same micro-occlusion model, including the same FITC-dextran, did not report such a phenomenon. We reason that this is due to the fact that histological analyses in these studies were performed at earlier time points following occlusion ([13, 25, 46]). Even when the pattern could have been observed, it went unnoticed. For example, Li and coworkers [19], used non targeted photothrombosis and studied the histological impact up to 14 days post stroke; their Figure 5, panel A6 (bottom) shows a clear Iba1-positive trail as reported in much more detail here. Indeed, we show that the trail is initially observed one week following occlusion and develops throughout time becoming more pronounced after 14 days (Figure 2), even for smaller cortical infarcts originated by the photothrombotic occlusion of a single penetrating vessel.

Our results point to microglia/macrophages as the key cellular components of the trail (Figure 5). As we did not observe neither dextran leakage from the periphery (Supplementary Figure S2), nor proliferation along the white matter (Supplementary Figure S7), nor differences in microglia density near the white matter (Supplementary Figure S8), we conclude that microglia/macrophages migrate from the infarct area along the white matter tracts. Microglia are the primary immune effector cells in the brain ([47]). Following injury, a cascade of pro- and anti-inflammatory cytokines and chemokines are released in the ipsilesional, as well as in the contralesional regions ([48]), resulting in microglial proliferation and migration. In aging ([49]) multiple sclerosis (MS) ([50]) and TBI ([48]), microglia/macrophages were shown to migrate and to accumulate in the white matter, where they cleared myelin debris from injured axons and allowed remyelination. Accordingly, we found that the microglia accumulation along the white matter was accompanied by myelin reduction in the heterozygous CX3CR1 mice but not in the knock-out (Supplementary Figure S10). Together, these observations emphasize the potential role of microglia to limit the detrimental impact of single and multiple microinfarcts.

We found that the microglial migration following micro-occlusion was fractalkine (FKN) CX3CR1 axis dependent (Figure 4). While infarct size (i.e. area and depth) was similar in CX3CR1-KO and 251 heterozygous animals, knock-out animals displayed shorter trails following occlusion of vessels with similar diameters. The FKN-CX3CR1 pathway underlies neuron-microglia communication in health and disease ([51]) and mediates microglial activation and migration ([38, 39]). This pathway is also key for microglial support of developing neurons and for synapse maturation and stability ([52]). In an MS model, CX3CR1-KO mice displayed reduced phagocytosis and microglial migration along the corpus callosum; resulting in inefficient clearance of myelin debris and defected remyelination ([50]). While we have not directly measured remyelination in the current study, lack of CX3CR1 resulted in altered reorganization 8 patterns throughout time, as indicated by DTI measurements (Figure 5). DTI is sensitive to structural cellular changes pointing to degenerative processes, including astrogliosis, de- and remyelination, and microglial/macrophage activation ([53, 54]). In addition, tractography analysis of the splenium section of the corpus callosum reveled local increase in FA values in CX3CR1-KOs and the distal increase in FA values in WT mice, indicating CX3CR1-mediated impacts on fiber integrity following micro-occlusions. While the anatomical basis for increased FA values is unknown, it may be explained by increased post 4 injury axonal sprouting, increased myelination, or gliosis ([55]). Notably, microglial activation along fiber tracts far from the infarct site was previously demonstrated using PET-MRI in stroke ([56]) and TBI ([57]) patients. In the current study we show that following single penetrating vessel occlusion microglia/macrophages migrate along the white matter tracts, accompanied by wide range structural reorganization, both of which were mediated by the fractalkine axis. In the current study we did not measure the functional impact of the microglial migration along the white matter tracts and of the observed reorganization patterns. However, taken together with the aforementioned studies regarding the role of microglia and the fractalkine axis in myelin debris clearance, remyelination and repair ([50] [58]), we hypothesize that following micro-occlusion, the cell bodies in the cortex are injured followed by axonal Wallerian degeneration ([59, 60]). Microglia/macrophages are then activated and migrate to clear myelin debris, allowing remyelination and repair. In cases of multiple microinfarcts, microglia/macrophages may not be effective enough or sufficient for repairing the damage, or may be over-activated due to a chronic condition [61, 62], resulting in neurodegeneration and cognitive impairments.

Many reports describe evidence for ipsilesional reorganization post stroke or photothrombotic occlusion (e.g. [63, 64]), and some findings point to contralateral reorganization post stroke [65, 66]. Van Meer and colleagues [67] found that in the first days after unilateral stroke in rats, interhemispheric functional connectivity disappeared, but were partially restored in the next few weeks, along with improved sensorimotor function [67]. Functional alterations are often accompanied by structural reorganization [68]; although in the current study we did not assess functional outcomes, the structural changes reported here might constitute the morphological basis for such post stroke functional changes. Moreover, since stroke results in diaschisis [69] (i.e. functional loss in an otherwise healthy brain region connected to an affected area) the microglia/macrophage migration described here could be a potential therapeutic target for preventing the development of such condition.

The growing evidence linking cortical microinfarcts with cognitive decline suggests that the tissue damage is not limited to the ischemic core and subcortical white matter (for a comprehensive review see [70]). Under this emerging concept, distant yet contiguous tissue, has been shown to be affected by microinfarcts [9]. The work we presented here radically redefine *distant* as encompassing both the ipsi- and contra-lesion hemispheres in a process that involves the likely degradation of axonal tracts. Moreover, we report here a new phenomenon associated with microinfarcts; namely the formation of a microglia/microphage migratory path, initiated at the microinfarct core which develops along the axonal mantle and reaches the contralateral hemisphere across the corpus callosum over a period of several weeks. Importantly, we showed that both the extent of remodeling and migration are modulated, at least partially, by the fractalkine receptor CX3CR1 expression levels. This later finding opens the opportunity for pharmacologically manipulation of microglia/macrophages activation to delay the detrimental impact of cortical microinfarcts.

## 4. Methods and Materials

### 4.1. Animals

All studies were approved by the Tel Aviv University ethics committees for animal use and welfare and in full compliance with IACUC guidelines. Two to four month old C57BL/6J, B6.129P-Cx3cr1tm1Litt/J heterozygous (i.e. CX3CR1^*GFP*/+^) and homozygous (CX3CR1^*GFP/GFP*^) (The Jackson Laboratory, stock 005582) and NG2-TdTomato male and female mice were used. The CX3CR1^*GFP*/+^) mice were generated by crossing C57BL/6J with (CX3CR1^*GFP/GFP*^) mice. NG2-TdTomato mice were used in the colocalization experiment (Figure 5). For that purpose, NG2-CreER transgenic mice (The Jackson Laboratory, stock 008538; [71]) were crossed with Rosa26-tdTomato (Ai14) reporter line (Jackson Laboratories, stock 007908). Four weeks old progeny received tamoxifen for 4 days, as described previously ([72]), to induce TdTomato fluorescent protein expression in NG2 cells. All animals were housed under standard vivarium conditions (221C, 12h light/dark cycle, with ad libitum food and water). No animals where excluded from the experimental groups once assigned. Although no specific randomization scheme was used while allocating animals to each group, no selection criteria was implemented and animals were randomly picked from their cages to enter into the different cohorts.

### 4.2. Two-photon imaging and photothrombotic occlusion

For vascular imaging and photothrombotic occlusion of single penetrating vessels, a polished and reinforced thin-skull (PoRTS) window was used, as previously described [18]. Briefly, Mice were anesthetized and maintained on 1.52% isoflurane in a 30/70 oxygen/N2O mixture, and body temperature was monitored and maintained at 37ΰC. Animals were fixed using a stereotaxic frame, and a 2.5mm^2^ of the skull was thinned to a width of 20-40m with a high-speed manual drill (Osada, drill bit #5) over the right somatosensory cortex (center point at 2.5mm lateral and 2.5mm posterior to bregma). The skull was cooled after each thinning cycle using artificial cerebral spinal fluid (ACSF) and special care was given to avoid breaking of the skull and provoking inflammation. The thinned region was covered with #0 thickness cover glass for optical access. A custom-made metal head frame was fixed to the contralateral skull using cyanoacrylate glue (Loctite 401) and dental cement (high-Q-bond, BJM labs) to enable head fixation during imaging. A 3D printed well was glued to the skull surrounding the window, to hold water while imaging with a water immersion lens. For cranial windows in MRI experiments, the metal head frame and plastic well were not used to avoid magnetic artifacts. Instead, these animals were fixed under the two-photon microscope using a stereotactic apparatus, and their skin was sutured over the window. Carprofen (0.05 mg/kg) was provided for analgesia. For two-photon imaging we used a modified microscope based on a Sutter MOM platform, (Sutter Inc) with a Leica 25X water immersion objective (NA 0.95). Linescan data was acquired using MPScope ([73]) and raster images with Scanimage ([74]) softwares. To image blood vessels and determine the vessel type (artery or vein) mice were injected with 5% fluorescent-dextran (Table 1) dissolved in ACSF (25l; retro-orbitally). C57BL/6J and NG2-TdTomato mice were injected with FITC-dextran (2000kDa) or Texas red (70kDa) and vessel type was determined by blood flow directionality using arbitrary linescans, as previously described ([75]). CX3CR1 mice were injected with Alexa Fluor 633 hydrazide (Invitrogen) for specific arterial labeling ([33]). For targeted single-vessel occlusion we used an approach previously described [24, 12, 76]. Mice were injected with Rose Bengal dissolved in saline (25l; retro-orbitally) and a green CW laser (532 nm; 0.1-0.5mW; Beta Electronics, Irvine) was immediately directed for 30 seconds to the lumen of the targeted vessel (12 to 25m in diameter) through the imaging objective. We assured the vessel was completely occluded for at least 30 seconds later to ensure the occlusion was not transient. Control mice underwent an identical procedure, including cranial window preparation, but were either deprived of Rose Bengal administration or laser illumination.

**Table 1:**
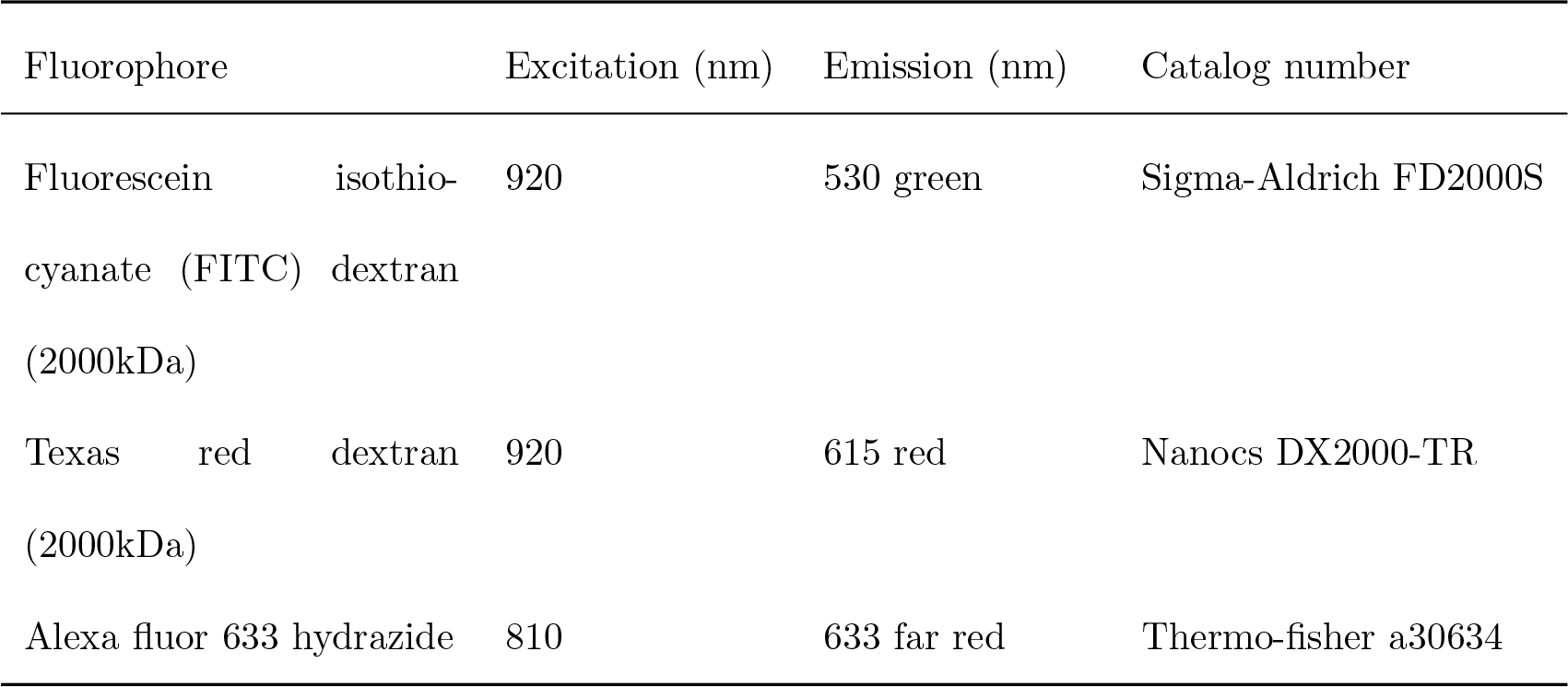
Fluorescent agents for blood labeling under Two-photon imaging.Description of agents used for all experiments involving vascular imaging under two-photon microscopy. Provided are the excitation wavelength used and the peak emission for each fluorophore.

### 4.3. BrdU labeling

For the proliferation assay, CX3CR1 heterozygous (n=6) mice were injected seven consecutive days (starting 2h after the occlusion) with thymidine analog 5-bromo-2-deoxyuridine (BrdU; 50mg/kg, i.p.; 7 Sigma-Aldrich, USA) as a marker for dividing cell. The mice were sacrificed seven days following the last injection (14 days following the occlusion).

### 4.4. Tissue processing and immunostaining

Following euthanization (Pentobarbital overdose), mice were transcardially perfused with PBS-heparin followed by 4% paraformaldehyde (PFA). Brains were postfixed at 4C for 24h, then cryoprotected in 30% sucrose for at least 48h. Coronal sections of 40m were cut using freezing microtome (Leica) and processed as free-floating brain sections. The brain sections were blocked for 1h in blocking solution (10% goat serum, 0.3% Triton-X in PBS) at room temperature and then incubated overnight at 4C with the following primary antibodies in blocking solution: Rabbit anti-Iba1 (1:500; Wako Chemicals); Goat anti-DCX (1:200; sc-8066, Santa Cruz); Rabbit anti-NeuN (1:1000;MABN140, Millipore); Rat anti-GFAP (1:1000; Zy130300, invitrogen); Mouse anti-MBP (1:500; SMI 99, BioLegend); Rat anti-CD45 (1:100, 550539, BD Bioscience); Rabbit anti-Olig2 (1:1000;AB9610, Millipore). For negative control, the primary antibodies were omitted and sections were incubated in blocking solution. The sections were incubated for 2h at room temperature with corresponding secondary antibodies: goat anti-rabbit 647 (1:1000; Invitrogen); Goat anti-mouse 647 (1:1000; Invitrogen); Goat anti-rat 647 (1:1000; Invitrogen); Donkey anti-goat 647 (1:1000; Invitrogen) in blocking solution, followed by DAPI (1:1000; 0215757405, MPbio) for nuclei visualization. To double stain with BrdU, before blocking, sections were incubated in 2M HCl for 20 minutes at room temp followed by neutralization in 0.1mol/L borate buffer and vigorous rinsing in PBS. The BrdU staining was performed with rat anti-BrdU (1:200, Abcam, Cambridge, UK). Fluorescence was detected with a Zeiss V12 fluorescent binocular (Zeiss, Germany) for entire coronal sections, and with Leica SP8 confocal microscope (SP8, Leica) for high resolution images using Leica Microscope Imaging Software. In order to characterize the identity of the cells that co-localized with FITC-dextran along the white matter, we counted the percent of cells on the trail that were double positive for FITC and the tested antibody. The absolute number of FITC-dextran labeled cells was counted in several fields of view along the corpus callosum using ImageJ software. Data represent the percentage of the cells that colocalized with FITC for each marker. All imaging and analysis parameters were kept constant, automated image processing and file naming was used as blinding approach.

### 4.5. Measurement of Infarct Area

For infarct area measurement, brains from perfusion-fixed animals were sectioned into 40m coronal 6 sections and stained for neuronal nuclei (NeuN). The central slice of the infarct was collected and imaged with Zeiss V12 fluorescent binocular (Zeiss, Germany). Infarct area of each slice was manually delineated using ImageJ software (NIH). Infarct area was calculated by measuring the area devoid of NeuN staining. For each animal the section with the largest infarct area was chosen.

**Table 2:** Antibodies used for staining. Description of anitbodies used for all immunohistochemistry stainings.

| Antigen | Host | Primary | Secondary |
| --- | --- | --- | --- |
| NeuN | Rabbit | 1:1000 MABN140, Millipore | Goat anti-rabbit |
| DCX | Goat | 1:1000; ab18723, Abcam | Donkey anti-goat |
| Iba1 | Rabbit | 1:500; 019-19741, Wako Chemicals | Goat anti-rabbit |
| GFAP | Rat | 1:1000; Zy130300, invitrogen | Donkey anti-rat |
| CD45 | Rat | 1:100, 550539, BD Bioscience | Donkey anti-rat |
| MBP | Mouse | 1:500; SMI 99, BioLegend | Goat anti-mouse |
| BrdU | Rat | 1:200, ab6326, Abcam | Donkey anti-rat |
| Olig2 | Rabbit | 1:1000; AB9610, Millipore | Goat anti-rabbit |

### 4.6. Measurement of Infarct depth

Infarct depth was calculated relatively to the cortex thickness in the area of infarct. The perpendicular distance from the pial surface to the upper border of the subcortical white matter was measured as total cortical thickness. Infarct depth was measured similarly, from the pial surface to the lower border of the infarct based on NeuN staining. Relative depth was calculated as the ratio between *infarct depth* and *cortical depth* (Figure 2 F). Based on this definition, if the infarct spanned the entire cortex and reached the white matter, the relative depth was 1.

### 4.7. Measurement of trail length

To quantify the trail length, the distance of positive FITC-dextran staining (in C57BL/J6 mice) or GFP (in CX3CR1 mice) was measured along the corpus callosum of the central infarct section. End points of the trails were manually determined along the corpus callosum using ImageJ software (NIH).

### 4.8. CX3CR1 density measurement along the corpus callosum

To quantify microglia/macrophage density along or above the corpus callosum in CX3CR1 mice, we 393 acquired whole brain coronal images using an Olympus slide scanner (Olympus IX83) in X20 magnification. To analyze these images, we designed an interactive Matlab GUI to mark the following components: the outline of the infarct area, the corpus callosum and five regions of interest (ROIs) along it. ROIs were selected as follows: two regions in the ipsilesional hemisphere, one in the midline and two in the contralesional hemisphere (Figure 5E). For contralateral hemisphere of photothrombotic occluded mice, the ROIs were selected in the approximate mirror regions of the ipsilesional ROIs. Similarly, the five ROIs for the density measurements near the corpus callosum were marked on the cortex just above the corpus callosum (Figure 5-Supplementary Figure S8). For control mice, cell density was assessed in the equivalent regions to the treated mice. For each of the marked ROIs, an auto-thresholding adaptive procedure was performed on the GFP channel to obtain a binary mask. Next, to remove thin cell processes and noise (in the form of smaller than cell soma binary blobs), the masks were initially eroded and a size restriction filter was subsequently applied. The specific filter parameters (erosion filter type and size and area threshold min and max values) are the only free parameters in the process and were chosen upon visual inspection of a sub set of images; once those were set they were used for all remaining images. The outcome of this process is an estimation of GFP densities in each ROI and the distance along the corpus callosum to the center of the infarct (projected to the corpus callosum). Data from several coronal slices (2-4) surrounding the infarct of each mouse brain were obtained from four-six mice of CX3CR1^GFP/+^ occluded or control animals.

### 4.9. Diffusion tensor imaging (DTI)

MRI scans were performed on a 7-Tesla MRI scanner (Bruker, Karlsruhe, Germany) with a 30-cm bore and a gradient strength of up to 400 mT/m. Volume coil for excitation and a mice quadrature coil for acquisition were used. Anesthesia was induced with 4% isoflurane (Vetmarket Ltd., Israel) and maintained with 12% isoflurane in 100% oxygen at a flow rate of 0.3-0.5 L/h. The respiratory rate was monitored throughout the entire experiment. The body temperature was monitored and maintained at 37C using a feedback system of circulating water. Wild type (n=9) and CX3CR1-KO (n=8) mice with MRI compatible thin skull windows were imaged before occlusion (i.e. baseline) and longitudinally at 1, 7, 21 and 28 days following micro-occlusion induction.

MRI protocol consisted of T2-weighted scans and diffusion tensor imaging (DTI). T2-weighted images were acquired with the following parameters: a MSME T2-weighted sequence allowed the acquisition of contiguous horizontal slices, which covered the whole mouse brain volume. Time of Repetition (TR) / Time to Echo (TE) =8115/14ms, FOV of 1.81.8cm^2^, matrix size of 192160, resulting in an in-plane resolution of 98118m^2^. The DTI images were acquired with the following parameters: Fast low-angle shot (FLASH), 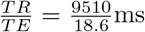,∆*/λ* = 10/4 ms,Δλ 32 non-collinear gradient directions with a single b-value at 1307 s/mm^2^ and an image of b-value of 0 s/mm2 (referred to as b0 images). FOV of 1.89 × 1.89 cm^2^, matrix size of 90 × 90 × 90 resulting in an in-plane resolution of 21m^3^. For both MRI protocols, 38 contiguous horizontal slices, 0.21mm thick, were collected. To improve signal to noise ratio (SNR), and compensate for head motion due to animals breathing, DTI acquisition time was 17 min, and the entire MRI protocol lasted 35 minutes for each session.

### 4.10. DTI analysis

MRI data was analyzed using ExploreDTI software ([77]). Prior to DTI calculation, the following preprocessing steps were performed: anisotropic smoothing, bias correction, skull stripping and regular-34 ization. DTI was then calculated and corrected for head motion and EPI distortions. Maps of diffusion weighted images (DWI), mean DWI and B-matrix data were exported. For voxel-based and region based analysis, mean diffusivity (MD) and fractional anisotropy (FA) were exported. Scans of all time points for each mouse were normalized into a standard space, an averaged FA image of 16 baseline scans, using a 12-parameter affine non-linear transformation in SPM8 (UCL, London). Following normalization, images were smoothed with a 0.4 mm Gaussian kernel.

### 4.11. Tractography analysis

To observe white matter reorganization following microinfarct, tractography procedure, which enables reconstruction of white matter fasciculi ([78]), was performed using ExploreDTI software ([77]). The dorsal part of the corpus callosum (splenium), that is located beneath the occlusion site, was used as seed region. Our fiber tracking parameters were 0.2 × 0.2 × 0.5 mm^3^ seed point resolution, 0.16mm step size and 300 angular threshold. The length of the fibers was transferred to mutual dimensions of total length of 100 sections, where the midline was at section fifty. FA values of each group, for every time point at each section, were averaged.

### 4.12. Statistical analysis

Statistical analyses were performed with MATLAB (MathWorks) or Prism (Graphpad) softwares. Linear regression was applied for analyzing correlations between trail length and time post occlusion, infarct area and relative depth. Gaussian based statistics (unpaired Student’s t-test) was applied, unless the assumptions were not met, there we used the nonparametric Mann-Whitney test. Data presented as meanSEM, n.s = not significant. In figure 5 (E) Mixed-effect General Linear Model (GLM) was used for GFP density calculation with group and distance from cyst as the independent variables. The model contained terms for the *intercept*, *group* (sham, occlusion), *distance* from infarct center (measured along white matter tract) and interaction between *group* and *distance*. Analysis was performed in Matlab using the fitlme function from the Statistical toolbox. In 1 (D-E), a voxel-based repeated measures ANOVA was performed within the WT group, with 5 time points as a within subject factor, to reveal tissue changes through time. One mouse was excluded from analysis due to extremely enlarged ventricles and general ill appearance. In figures 5 and 6, a voxel-based repeated measures ANOVA with main effects between groups (WT and KO) and within time (5 time point) and interaction (group vs. time point) was obtained as statistical parametric maps, followed by multiple comparison correction (FDR). A value of p<0.05 was considered statistically significant. For illustration purposes we show the clusters both before and after FDR correction. The parametric maps are superimposed on an averaged FA map of baseline scan of all mice. For the tractography analysis, ANOVA was performed on the average FA value of the region (ipsilesional (1-49) and contralesional (51-100) sections), between two time points (baseline and 4 weeks after the occlusion). Tukey post-hoc analysis on each section was performed to find the sections in which there was a significant difference between the means of the two time points. The specific test can be found in the results section.

## 5. Acknowledgments

This research was funded by the FP7-CIG Marie Curie Action to PB (project 618251); PB would also like to thank the support from the Israeli Science Foundation (1019/15) and the Leducq Foundation (FDNLEDQ-15CVD-02). AL thanks the Fondation Judaism Français for doctoral scholarship support. AB is grateful to Dante family for the award of doctoral fellowship. The authors thanks Amos Gedalyahu for comments on earlier version of the manuscript.

## 6. Author Contribution

Conceptualization, AL, AB and PB. Methodology, AL, AB, YA and PB, Software HH and PB. Investigation, AL, AB, HH and HL.Resources, PB. Writing original draft, AL, AB and PB. Writing Review & Editing AL,AB, YA and PB. Visualization, AL,AB, YA and PB. Supervision YA and PB.

---

**Supplementary Fig S1:**
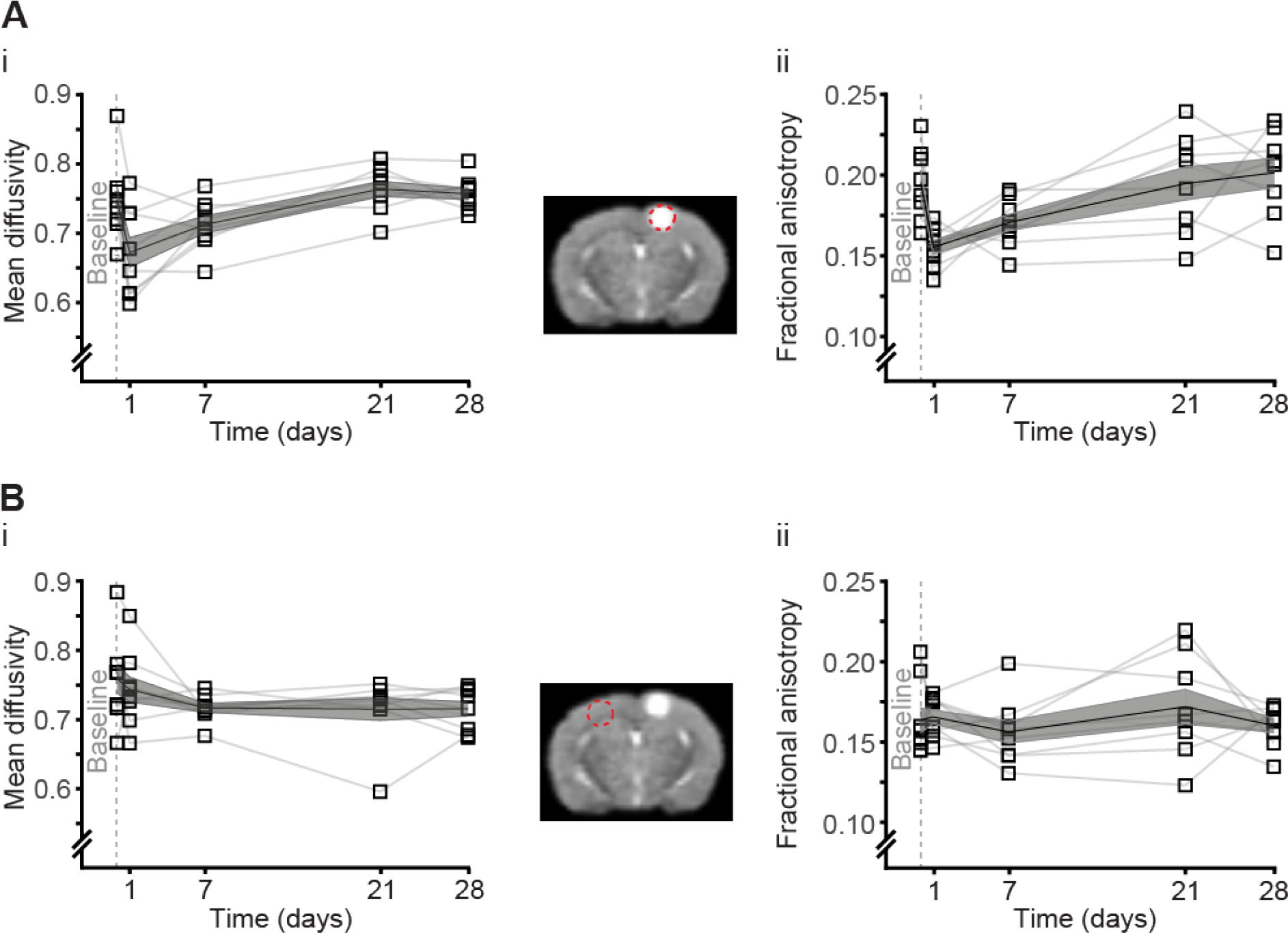
Region based analysis of the occlusion and contralateral area (A) Region-based analysis of MD (i) and FA (ii) values in the occlusion area somatosensory (SS) cortex show tissue alterations in response to the micro-infarct, while (B) presents the mirror region in the contralateral hemisphere where the MD (i) and FA (ii) values do not change through time in response to the occlusion.

**Supplementary Fig S2:**
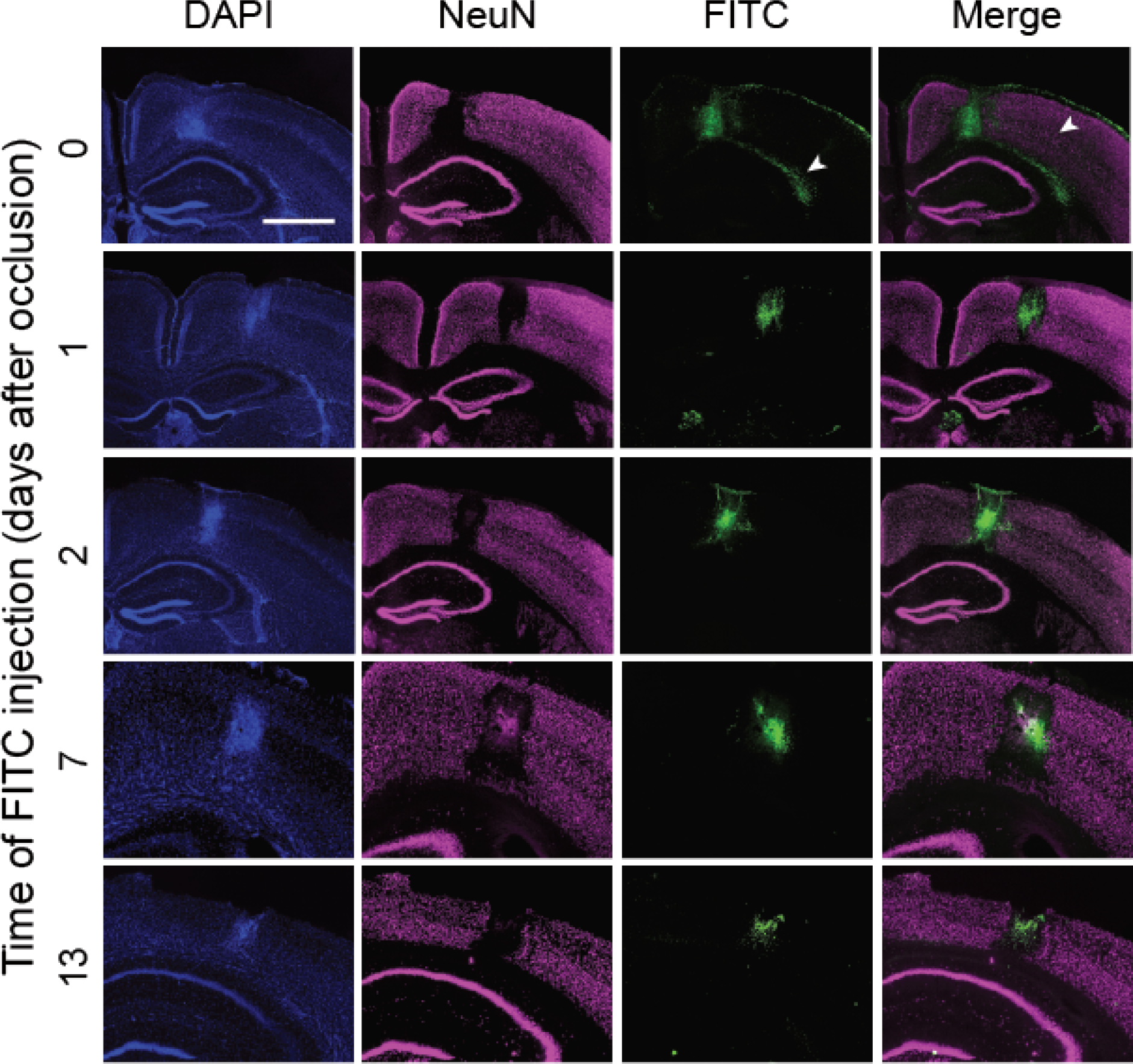
FITC did not infiltrate to the white matter from the periphery. FITC injection to the bloodstream at 0,1,2,7 and 14 days post occlusion with Texas Red (2000 kD) shows a presence of the FITC-dextran on the corpus callosum only on day 0 but none at later time points. Scale bar: 1mm.

**Supplementary Fig S3:**
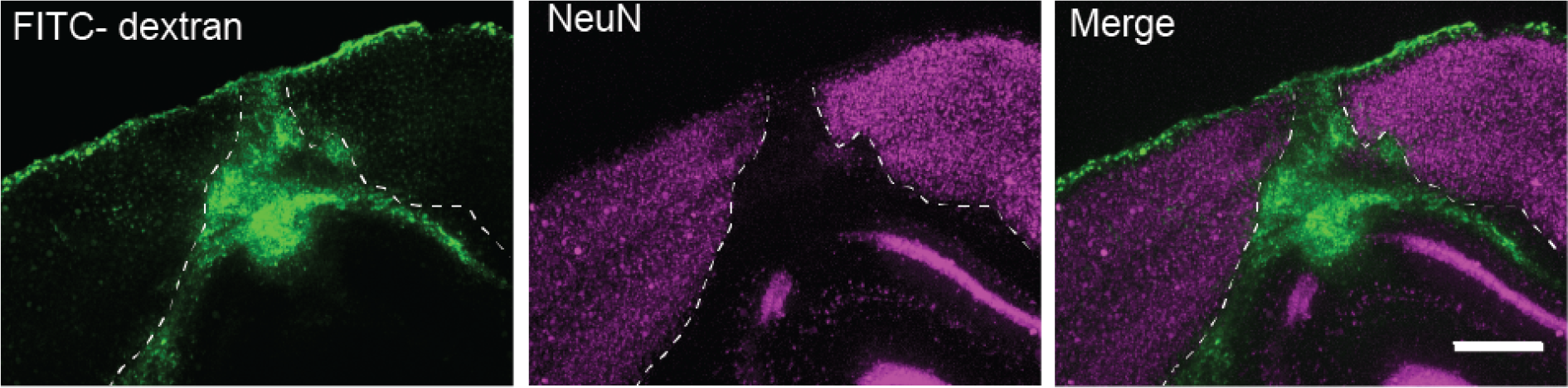
FITC-dextran accumulation was observed in the pial surface. Scale: 500m

**Supplementary Fig S4:**
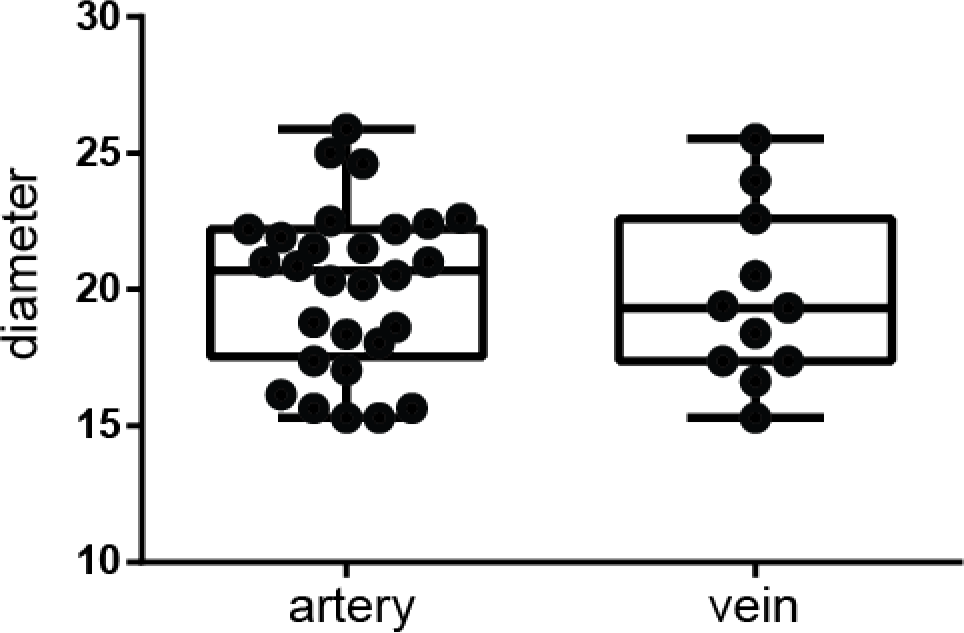
Distribution of the diameter of occluded blood vessels in wild type mice. The average diameter between the genotypes was similar (20.10.56m for arteries n=28 and 19.670.96m for veins n=11; p=0.707, Students t-test)

**Supplementary Fig S5:**
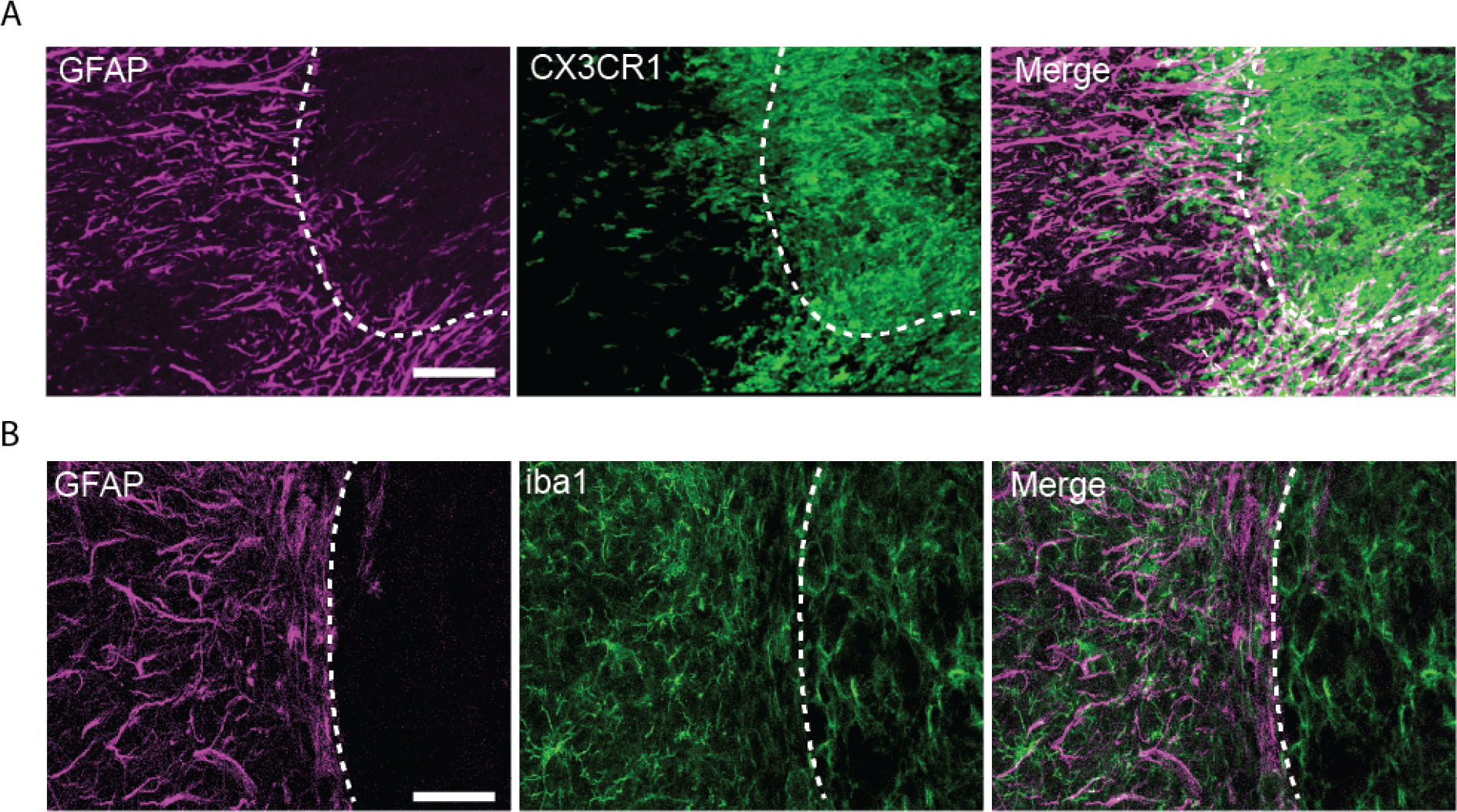
Astrocytes surrounded the injured site and formed glial scar, but did not enter the lesion core. High-magnification of the infarct border where the astrocytes create glial scar surrounding the core. In the ischemic cortex, a clear distinction is seen between the astrocytes (GFAP, magenta) outside the infarct area and the microglia/macrophages that are inside the core, in (A) CX3CR1-GFP mice and (B) WT mice with iba1 staining (green). Scale: 20m.

**Supplementary Fig S6:**
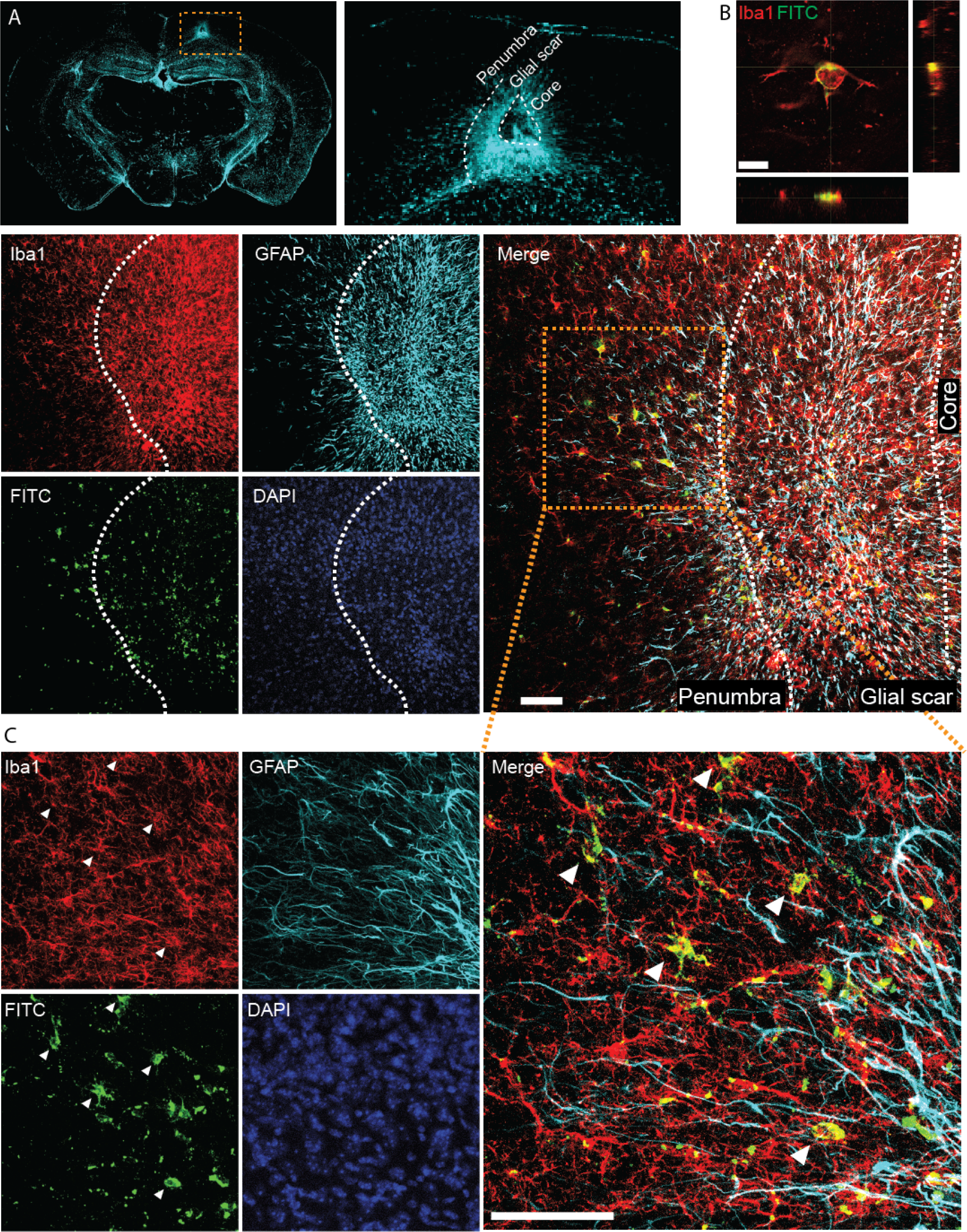
FITC colocalized with IBA1 but not GFAP at the penumbra. (A) Magnification of the glial scar in the infarcted cortex. (B) The FITC-dextran (green) that leaked to the cortex was colocalized with microglia/macrophages (red). there was no double labeling of FITC with astrocytes (cyan) (white arrowheads point to the colocalized cells). White dashed line demonstrates the border between the glial scar and the penumbra. (C) Enlargement of orange dashed box in A. Scale bar: A: 50m, B: 10m, C: 50m.

**Supplementary Fig S7:**
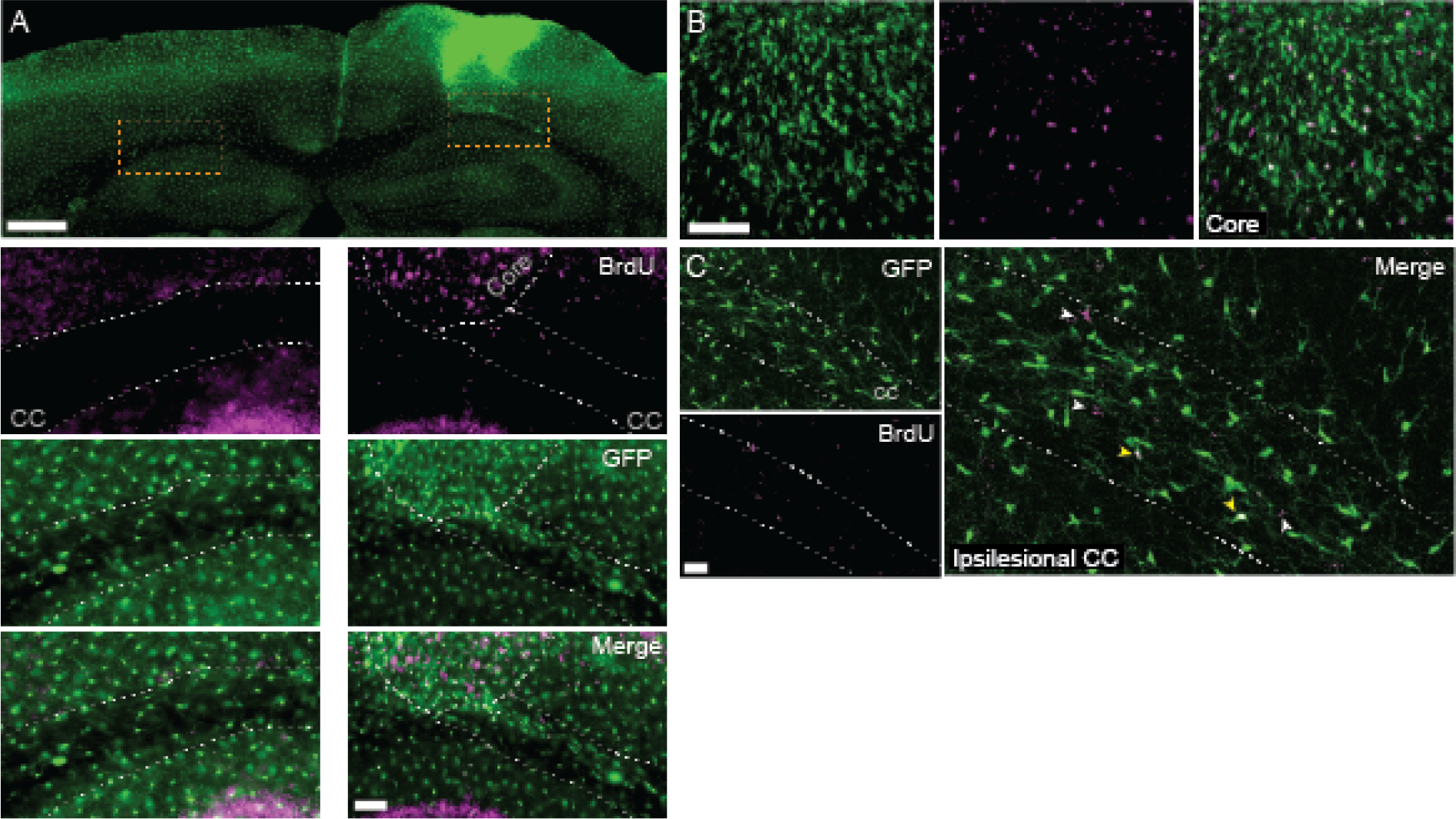
Only a small amount of the GFP+ on the cortex callosum are newly generated microglia/macrophages. (A) BrdU double labeling for newly formed cells and microglia/macrophages in CX3CR1^GFP/+^ mice, in the ipsi- and contralesional hemispheres. Only a small number of cells were double labeled for BrdU (magenta) and GFP (green). (B) magnification of the core where many cells are GFP+/BrdU+. (C) magnification of the ipsilesional corpus callosum. white arrowheads point to BrdU labeled cells and yellow arrowhead points to GFP/BrdU double labeled cells. Scale bar: A:1mm, 50m, B:50m, C:100m

**Supplementary Fig S8:**
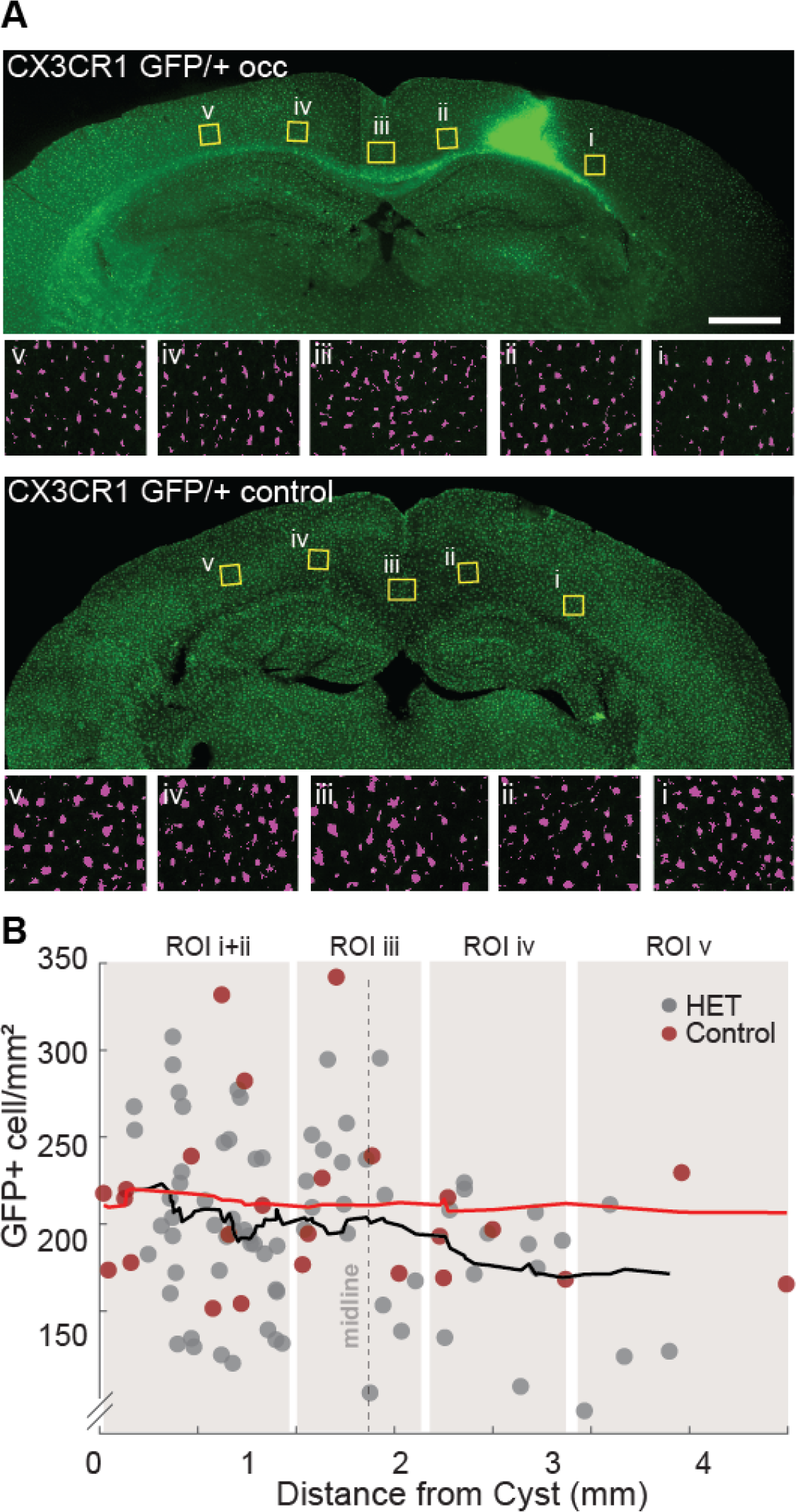
GFP density above the corpus callosum did not change. (A) GFP+ density was measured in 5 ROIs above the corpus callosum of CX3CR1^GFP/+^ occluded and control mice, 14 days after micro-occlusion induction. Representative images of automatically identified cells in each ROI are shown below each group (magenta). (B) Mixed-effect general linear model showed no significant effect for any of the model terms but the intercept (p<0.0001, DF = 96). Scale bar: 1mm.

**Supplementary Fig S9:**
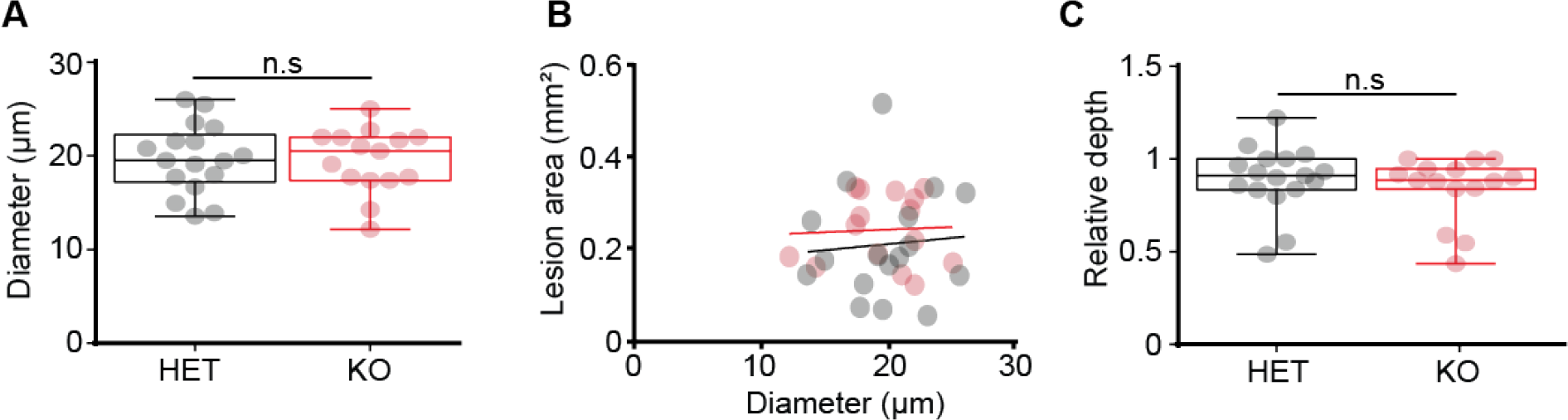
Properties of occluded arteries of CX3CR1 heterozygous and KO mice. (A) The average occluded arteries diameter between the genotypes was similar (19.70.89mm for HET n=17 and 19.510.88 mm for KO; p=0.788, Students t-test). (B) No correlation between occluded artery diameter and infarct area was found for HET (R^2^=0.006, p=0.755, black) and KO (R^2^=0.002, p=0.854, red) mice. (C) No difference in infarct depth for both genotypes (0.890.04 for HET n=17 and 0.840.04 for KO; p=0.39, Students t-test).

**Supplementary Fig S10:**
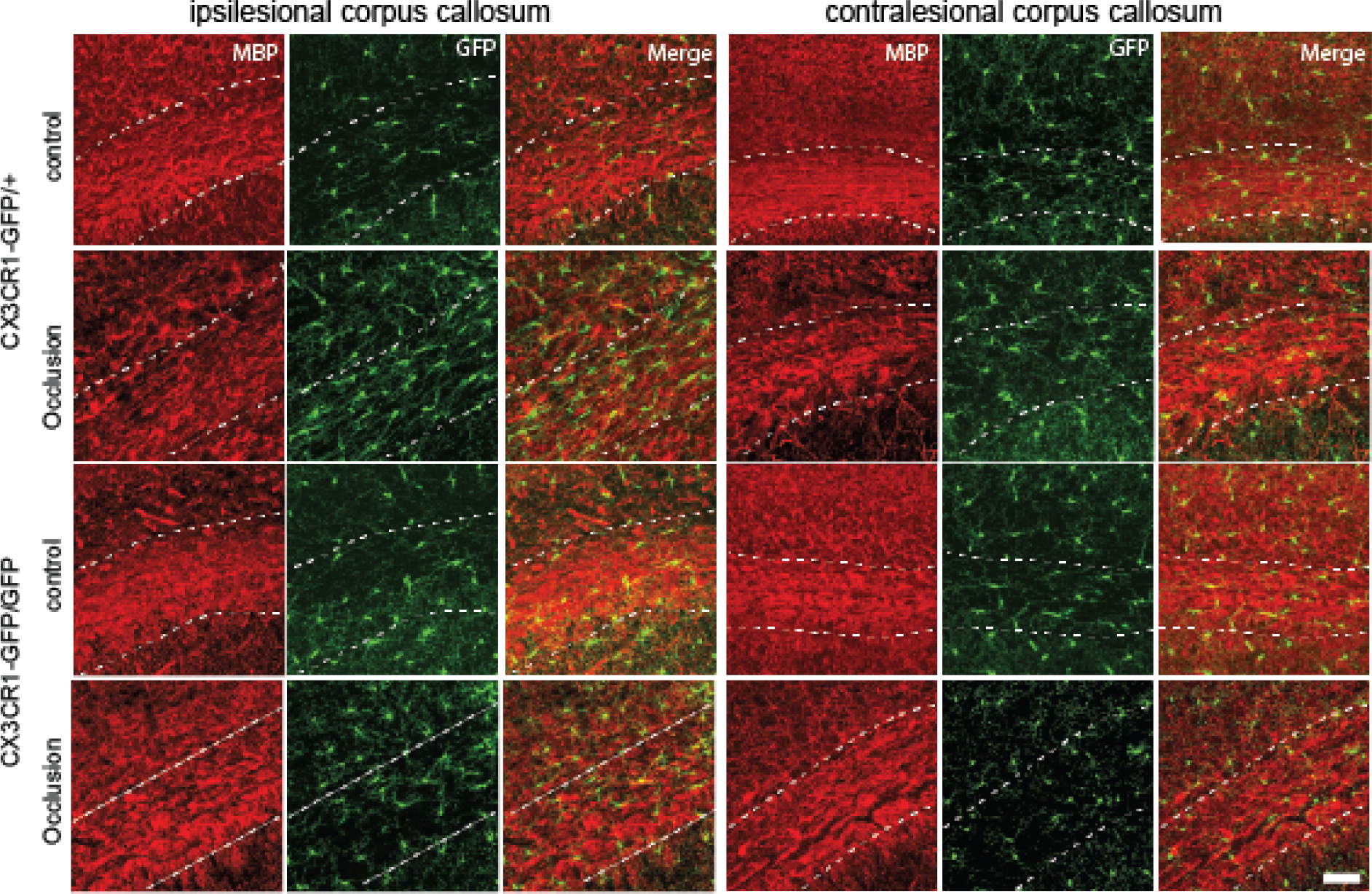
Immunofluorescence labeling of myelin basic protein (MBP) in CX3CR1^GFP/+^ and CX3CR1^GFP/GFP^ mice 14 days following photothrombotic occlusion. In control animals the myelin appears intact (red) in both genotypes, while following occlusion ipsilesional structural aberrations appeared in the corpus callosum of heterozygous mice along with increase in CX3CR1 density (green). These changes in myelin integrity were milder in KO mice and did not reach the contralateral hemisphere. CX3CR1-positive cells are denser and have ramified morphology with processes extending along the myelin fibers. Scale bar: 100m

